# Gene-based calibration of high-throughput functional assays for clinical variant classification

**DOI:** 10.1101/2025.04.29.651326

**Authors:** Daniel Zeiberg, Ross Stewart, Shantanu Jain, Malvika Tejura, Abbye E. McEwen, Shawn Fayer, Yuriy Sverchkov, Mark Craven, Vikas Pejaver, Alan F. Rubin, Lea M. Starita, Douglas M. Fowler, Anne O’Donnell-Luria, Predrag Radivojac

## Abstract

High-throughput assays measure a broad range of variant effects on gene function and hold promise for supporting genomic medicine. Current clinical guidelines for rare Mendelian diseases rely on establishing gene-specific score thresholds for each assay that separate pathogenic from benign variants. This introduces inconsistencies and subjectivity, ultimately lacking the rigor of calibration; i.e., mapping a variant score to a probability of pathogenicity. To address this problem, we introduce a semi-supervised framework for calibrating experimental assay data and propose Experimental score CALIBRator (ExCALIBR), a method that jointly models pathogenic, benign, synonymous, and population variants using skew normal mixtures to produce variant-specific probabilities of pathogenicity. Evaluated across 80 datasets from 39 genes, all meeting fit quality criteria, ExCALIBR substantially outperformed existing field standards and was further validated on the All of Us biobank data. Our results demonstrate that calibrated experimental assays generate indispensable evidence that will dramatically reduce variants of uncertain significance.

## Introduction

The American College of Medical Genetics and Genomics (ACMG) and the Association for Molecular Pathology (AMP) maintain evidence-based guidelines for classifying variants in genes and their use in genomic medicine [1]. Under this framework, variants are assigned to one of five categories based on their probability of causing disease as pathogenic (≥99%), likely pathogenic (≥90%), likely benign (*≤*10%), benign (*≤*1%), and variant of uncertain significance (10–90%). Classification depends on available evidence supporting or refuting disease causality. The guidelines specify the type (e.g., experimental, computational, population, or segregation data), strength (supporting, moderate, strong, very strong, standalone), and direction (benign, pathogenic) of evidence, along with rules for combining evidence to reach a final classification [1]. Pathogenic/likely pathogenic (P/LP) variants are considered diagnostic and may be clinically actionable, whereas benign/likely benign (B/LB) variants can be excluded from consideration. Variants of uncertain significance (VUS), which comprise a large majority of variants in ClinVar [2, 3] and are returned on one third of clinical testing reports [4], should not be used for medical decision making [5].

The ACMG/AMP system has been modeled by a Bayesian point-based system [6, 7] that improves rigor, but requires methodology for calibrating individual lines of evidence. For each evidence source, calibration aims to map a quantitative score (e.g., computational prediction, casecontrol counts, experimental assay readout) to a posterior probability of pathogenicity, which is then converted into discrete evidence of a specific direction and strength to support final classification [1, 7]. Calculating this posterior probability requires estimation of the positive likelihood ratio at a variant’s observed score and the variant’s prior probability of pathogenicity in a reference set. Proper calibration ensures that at each observed score, the expected fraction of pathogenic variants in a reference population equals the posterior probability computed at that score.

Experimental assays, including multiplexed assays of variant effect, are a scalable source of evidence for reclassifying VUS [8, 9] and can be prospectively deployed to assess the functional impact of variants not yet observed in the clinic [3, 10, 11]. The ClinGen Sequence Variant Interpretation Working Group developed recommendations for evaluating experimental evidence, establishing a framework for assessing assay validity and determining appropriate evidence strength based on the separation of pathogenic and benign control variants in the assay [12]. Assay score intervals that correspond to functionally normal and functionally abnormal variants are defined by the study authors using a range of approaches, including by visual inspection of score distributions. Positive likelihood ratios estimated for each assay interval are then combined with a prior probability of pathogenicity, defined by expert opinion, to assign uniform evidence strength to all variants within each interval. This approach, however, is subjective and incorrectly estimates the strength of experimental evidence for variants scoring near and far from the thresholds (Figure 1). Moreover, current approaches do not take advantage of internal controls such as synonymous variants to assess assay quality and variability [13]. This motivates the development of automated methods that calibrate experimental assay scores at the variant level by computing variant-specific positive likelihood ratios, estimating a prior probability of pathogenicity empirically from data rather than by expert opinion, and exploiting the distribution of synonymous variants.

**Figure 1.**
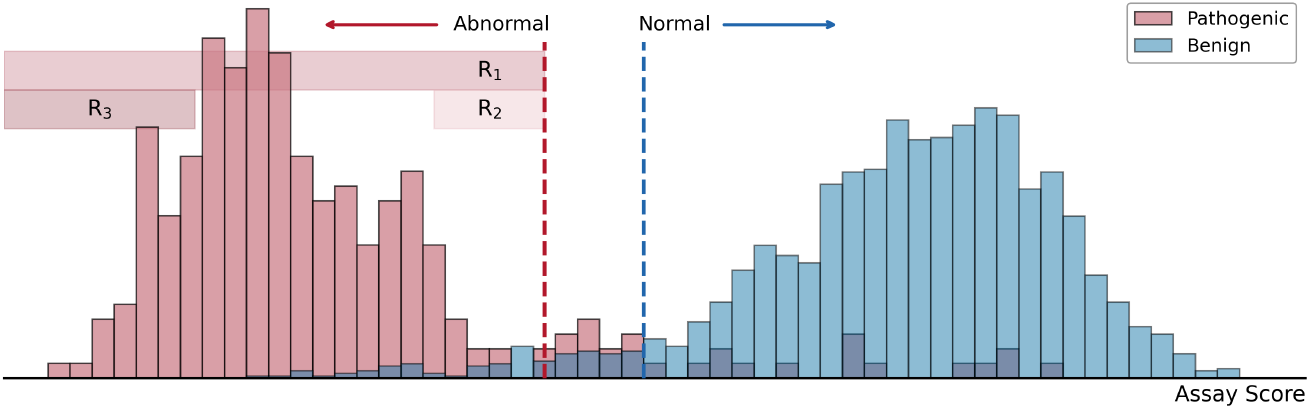
Illustrating the current use of experimental data in clinical variant classification. Current ClinGen recommendations rely on thresholds defining functionally normal and abnormal variants given a set of experimental scores. The evidence strength for variants exceeding these thresholds is computed using pathogenic (red) and benign (blue) control sets. The pathogenic evidence strength is computed as the ratio of true positive rate and false positive rate in region R_1_. Since experimental data are continuous, rather than binary functional labels, assigning evidence strengths at the variant level can result in more accurate use of experimental evidence. Variants with scores in region R_3_, far below the threshold, should be assigned higher evidence strengths than those assigned to variants with scores in region R_2_, slightly below the threshold.

Here, we present a new semi-supervised framework for calibrating experimental data and introduce Experimental score CALIBRator (ExCALIBR), a statistical method that jointly models assay score distributions of pathogenic, benign, population, and synonymous variants as skew normal mixtures. ExCALIBR estimates a variant’s prior probability of pathogenicity from a reference population sample, enabling posterior probability calculation and robust calibration despite the limited number of pathogenic and benign variants for many genes. We evaluate this method on a curated set of 80 experimental datasets that quantify the functional effect of variants in 39 clinically relevant genes, approaching 1% of disease-associated genes [14]. Our results show that ExCALIBR provides a good fit across all datasets, assigns variant-level evidence strengths that improve discriminatory power over conventional threshold-based approaches (accuracy: 97.9% vs. 93.6%), and successfully calibrates an additional 34 assays that could not be evaluated following the ClinGen-specified framework [12]. We validate our evidence assignments through gene-disease associations in the All of Us biobank [15], where 14 of 17 gene-disease pairs showed significant associations between pathogenic evidence strength and disease phenotypes. ExCALIBR not only allows new insights into experimental evidence as a whole, but also represents a fundamental shift from expert-defined thresholds and priors to empirical, unbiased evidence assignment that scales with rapid experimental data production [16–18].

## Results

ExCALIBR is a method that meets the requirements of *sensu stricto* calibration by estimating the posterior probability of pathogenicity for each assayed variant individually. The method jointly models score distributions of known P/LP and B/LB variants together with population and synonymous variants using mixtures of skew normal distributions with shared parameters. ExCALIBR estimates the prior probability of pathogenicity using gnomAD [19] as a population reference and computes local (variant-specific) positive likelihood ratios from modeled ClinVar [2] pathogenic and aggregated benign and synonymous mixture densities at a variant’s score. These quantities yield the posterior probability of pathogenicity, which is then translated into evidence points (±1, ±2, ±4, ±8) compatible with the ACMG/AMP guidelines, though our framework supports all integer evidence strengths between −8 and +8 and is extendable to real-valued evidence strength. ExCALIBR assigns robust, variant-specific evidence by performing 1,000 bootstrap iterations [20] per dataset and requiring that at least 95% of bootstrap models reach a given evidence strength at the variant’s score. For the analyses presented here, unless otherwise stated, evidence was assigned in an out-of-bag manner by aggregating assignments from bootstrap models that excluded the variant during fitting, ensuring unbiased evaluation. We applied ExCALIBR to 80 datasets across 39 clinically relevant genes from a harmonized resource recently developed by the Impact of Genomic Variation on Function (IGVF) Consortium [21]; see Methods.

### ExCALIBR fits diverse score distributions and reflects assay scope

To demonstrate the utility of our method, we visualized the model fits and score thresholds for four datasets in Figure 2. The models fit the score distributions well, demonstrating the flexibility of skew normal mixtures for modeling experimental data. Figure 2A shows that ExCALIBR assigns pathogenic and benign evidence to experimental datasets in which the assay separates the P/LP and B/LB distributions. In contrast, Figure 2B demonstrates that the model appropriately does not assign benign evidence to variants with mechanisms of pathogenicity not captured by the particular assay. This reflects limitations of the abundance assay, which primarily assesses the impact on protein stability and does not measure all types of functional impacts such as enzymatic activity [26], thus leaving a significant proportion of pathogenic variants with scores similar to those of synonymous variants [23]. Furthermore, Figure 2D illustrates that ExCALIBR does not assign evidence when the assay does not sufficiently separate the P/LP and B/LB distributions. This reflects what we intuitively understand about the strengths and limitations of these assays. Evidence strength thresholds for all integer levels between −8 and +8 and model fits for each dataset are available in are available in Supplementary Data 1–2.

**Figure 2.**
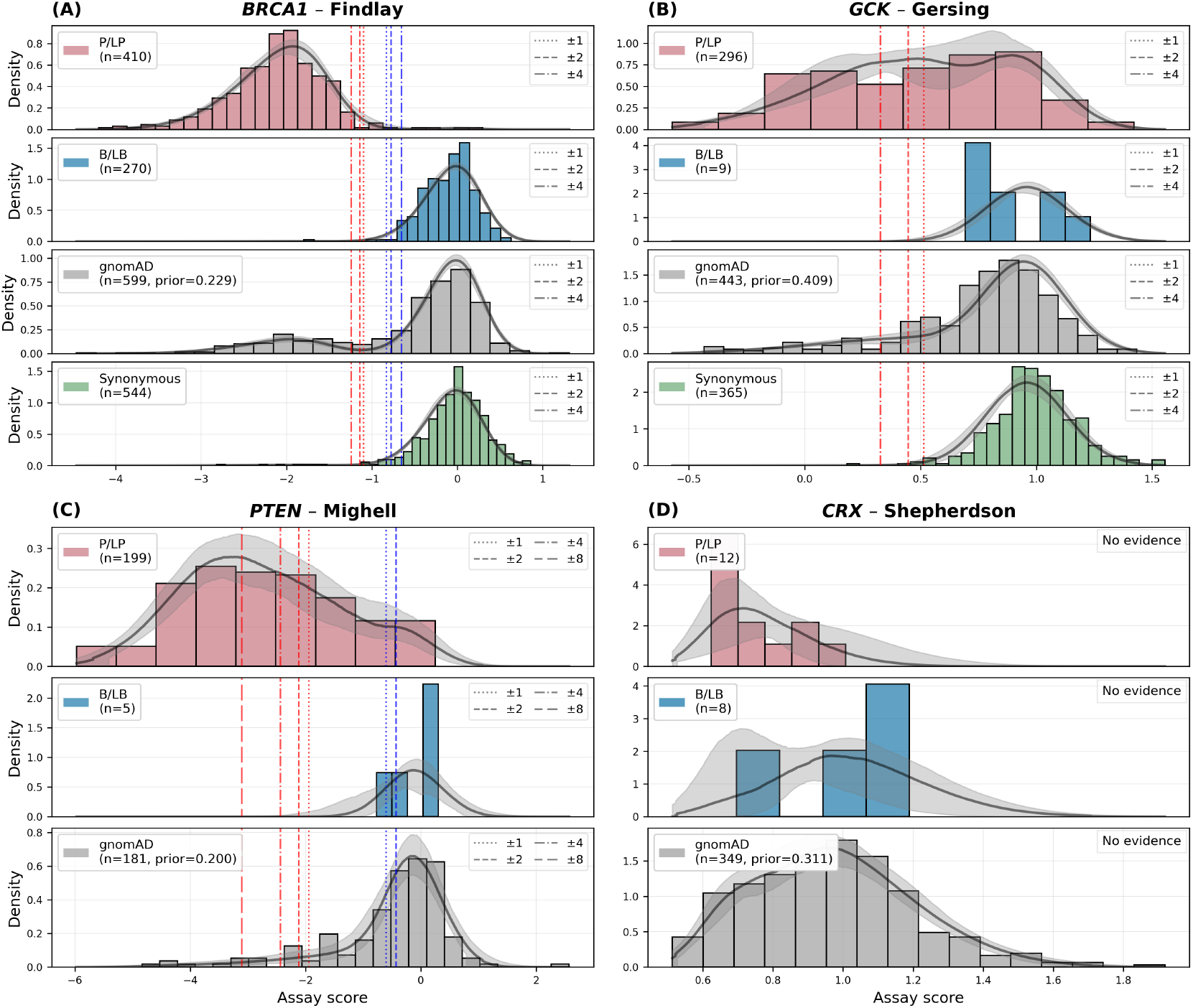
Example model fits and score thresholds for four experimental datasets assessing the effect of *BRCA1* variants (panel A), *GCK* variants (panel B), *PTEN* variants (panel C), and *CRX* variants (panel D). Datasets for panels A, B, C, and D were taken from Findlay *et al*. [22], Gersing *et al*. [23], Mighell *et al*. [24], and Shepherdson *et al*. [25], respectively. Median density across all bootstrap iterations is displayed along with the estimated 95% confidence interval. Score thresholds corresponding to each pathogenic and benign evidence strength are plotted in red and blue, respectively. Score thresholds corresponding to intermediate evidence strengths (±3, ±5, ±6, ±7) are excluded from visualization.

To quantify model fit quality, for each bootstrap iteration of a given assay we computed the normalized Yang *et al*. [27] distance between the empirical and fitted cumulative distribution functions for each sample. This distance ranges from 0 to 1, where lower values indicate better fit; see Methods for details. We considered distances below 0.2 to indicate a good fit. The gnomAD and synonymous samples are the most reliable indicators of fit quality due to their larger sample sizes; all 80 datasets achieved distances below 0.2 for these samples. Due to limited sample sizes and potential ascertainment bias, distances for P/LP and B/LB samples are generally larger and less reliable for assessing fit quality; nonetheless, 78 datasets (98%) met this threshold across all available samples (Supplementary Data 1). The two datasets (from *CARD11* and *SCN5A*) that exceeded this threshold failed to sufficiently fit the P/LP sample, where very few P/LP variants were available for fitting (*n <* 5). Supplementary Figure S1 shows the distribution of distances for the four datasets in Figure 2 across the 1,000 bootstrap iterations for each sample. Together, these results demonstrate that our method reliably fits diverse score distributions and assigns evidence consistent with known properties of these assays.

### Evidence distributions are concordant with functional annotations and ClinVar classifications

We compared the distributions of variant-specific evidence strengths assigned by our method to the functional annotations provided by the study authors across 39 datasets in 28 genes (Supplementary Data 1), excluding meta-analyses and datasets without author annotations (Figure 3A). To mitigate circularity, we excluded ClinVar controls deposited after 2018 for genes with published experimental assays that may have contributed to ClinVar classifications (*BRCA1, MSH2, PTEN*, and *TP53* ).

**Figure 3.**
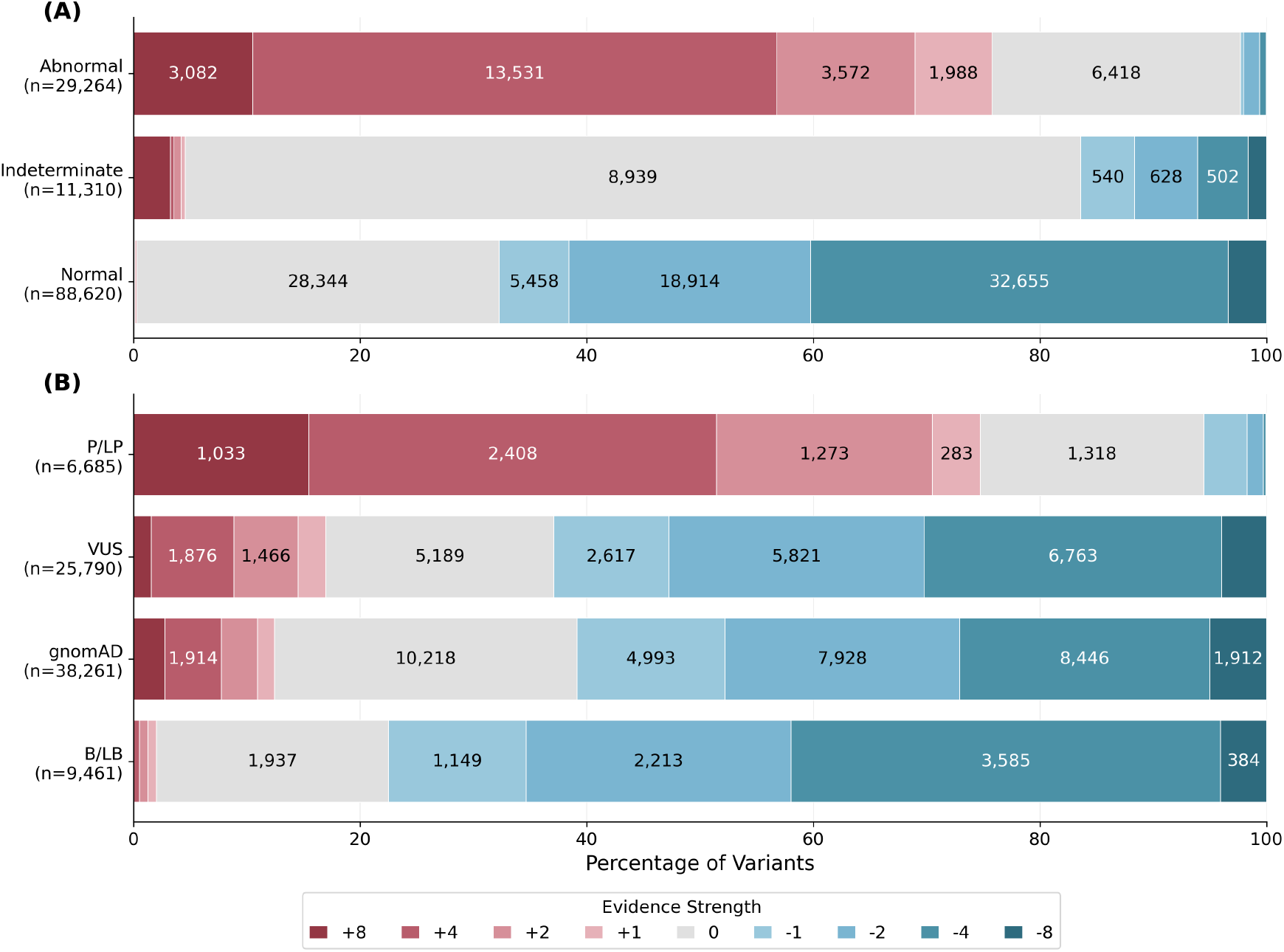
(A) Distribution of out-of-bag evidence strengths assigned by ExCALIBR for all variants with functional annotations provided by the authors of experimental studies (datasets provided in Supplementary Data 1). Author annotations were harmonized to a consistent vocabulary. (B) Distribution of out-of-bag evidence strengths assigned by ExCALIBR for variants in the P/LP, VUS, gnomAD, and B/LB samples. Each dataset was treated independently, and variants of all types (e.g., missense and synonymous) were included in the comparison, although synonymous variants were modeled as a distinct sample in ExCALIBR. Intermediate evidence strengths were floored to standard levels (±1, ±2, ±4, ±8) for visualization.

Of the variants annotated by the study authors as functionally abnormal, 76% were assigned pathogenic evidence of some strength, and 68% of variants annotated as functionally normal were assigned benign evidence of some strength. These reductions reflect that fixed categorical thresholds do not account for varying evidence strength across scores; variants near thresholds that authors categorize as functionally abnormal or normal may not reach sufficient evidence thresholds in our calibration framework, resulting in indeterminate evidence assignment. Fifty-seven percent of variants annotated as functionally abnormal by authors reached pathogenic evidence of strong (+4), and 11% of variants annotated as functionally abnormal reached pathogenic evidence of very strong (+8) using the model. Detailed gene-level statistics are presented in Supplementary Table S1.

Next, we compare the evidence strength distributions for variants present in ClinVar and gnomAD across all 80 datasets (Figure 3B). Of the 6,685 P/LP variants, 75% were assigned pathogenic evidence strengths and 20% received indeterminate assignments; only 6% were assigned benign evidence strengths. Of the 9,461 B/LB variants, 78% were assigned benign evidence strengths and 20% were indeterminate; only 2% were assigned pathogenic evidence strengths. Eighty percent of the 25,790 VUS were assigned either pathogenic or benign evidence strengths, with 63% assigned benign evidence strengths and 17% assigned pathogenic evidence strengths, suggesting the need for reclassification of ClinVar variants using updated experimental evidence. Comprehensive gene-level statistics are shown in Supplementary Table S2. As a whole, these results show that ExCALIBR evidence assignments are broadly concordant with both functional annotations and ClinVar classifications and provide quantitative evidence strengths that support reclassification of a substantial fraction of VUS.

### ExCALIBR improves accuracy and can calibrate more datasets

To demonstrate the impact that ExCALIBR could have on variant classification, we categorized our variant-level evidence assignments for P/LP and B/LB variants based on the direction of evidence and compared these to the author-provided functional annotations for the same 39 datasets described in the previous section. Author-provided assignments were given benefit of the doubt since some of the P/LP and B/LB variants from ClinVar may have been used to decide on the thresholds provided in their studies, whereas our evidence was evaluated using a strict out-of-bag approach. Figure 4 shows the number of P/LP and B/LB variants that our model assigned pathogenic or benign evidence directionality versus the number assigned by each author-provided functional class.

**Figure 4.**
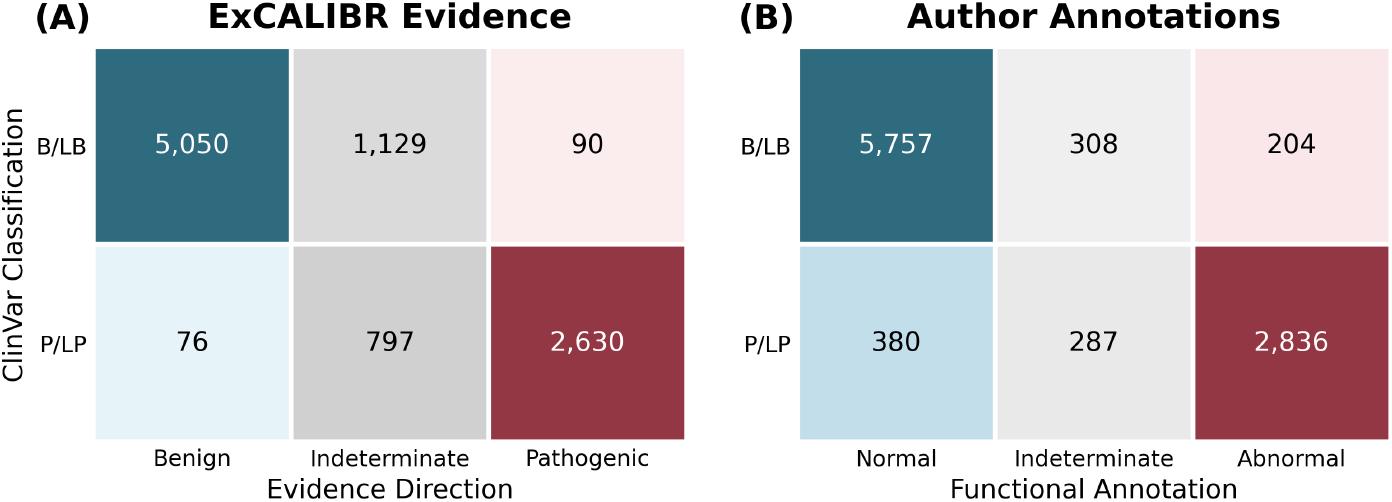
Comparison of ExCALIBR out-of-bag evidence assignments (A) and author-provided annotations (B) for ClinVar P/LP and B/LB variants in author-annotated datasets. Evidence assignments less than zero indicate benign direction; greater than zero indicate pathogenic direction; zero is indeterminate.

Categorizing author-determined functionally abnormal and normal variant assignments as pathogenic and benign, respectively, and P/LP and B/LB variants serving as positive and negative controls, we computed accuracy, sensitivity, specificity, Matthews correlation coefficient (MCC), global positive likelihood ratio (LR^+^), and diagnostic odds ratio (DOR); see Supplementary Table S3. Variants assigned indeterminate evidence were excluded from accuracy assessments. Our variant-level assignments considerably outperformed author-provided annotations (DOR: 1941.7 vs. 210.6) across all metrics at the cost of lower determinate assignment rates (80.3% vs. 93.9%). To include indeterminate assignments in assessment, we also calculated LR^+^ and DOR by dichotomizing the three-way classifications (Supplementary Table S3). For pathogenic evidence, we classified variants as pathogenic vs. non-pathogenic (benign or indeterminate), yielding LR^+^ = 52.3 and DOR = 206.8 for our method, compared to LR^+^ = 24.9 and DOR = 126.4 for author annotations (abnormal vs. normal or indeterminate). For benign evidence, we classified variants as benign vs. non-benign (pathogenic or indeterminate), yielding LR^+^ = 37.1 and DOR = 186.8 for our method, compared to LR^+^ = 8.5 and DOR = 92.4 for author annotations (normal vs. abnormal or indeterminate).

Figure 5A illustrates the accuracy comparison across 28 genes; two genes with all indeterminate ExCALIBR assignments were excluded. Of the remaining 26 genes, 22 (84.6%) showed improved accuracy with ExCALIBR evidence assignment. Next, we compared how many datasets reached each evidence strength for ExCALIBR versus the approach that derives evidence strengths from author-provided categorical annotations, as described in Brnich *et al*. [12]. Figure 5B shows that our method is able to assign evidence to substantially more datasets across each evidence strength, owing in part to ExCALIBR calibrating an additional 34 datasets that could not be evaluated with the Brnich *et al*. [12] approach (Supplementary Data 3). Finally, we validated our evidence strength assignments through gene-disease associations from the All of Us biobank [15]; see Methods for details. Figure 5C displays the odds ratios for the occurrence of disease in individuals with variants meeting each evidence strength for six example genes (*BAP1, BRCA1, MSH2, RAD51D, TSC2*, and *KCNQ4* ). For this analysis, all bootstrap iterations were used to determine evidence strengths; no filtering was performed to ensure out-of-bag evidence assignment. Fourteen of 17 genedisease pairs had statistically significant associations of pathogenic evidence and disease phenotypes (Supplementary Figure S2).

**Figure 5.**
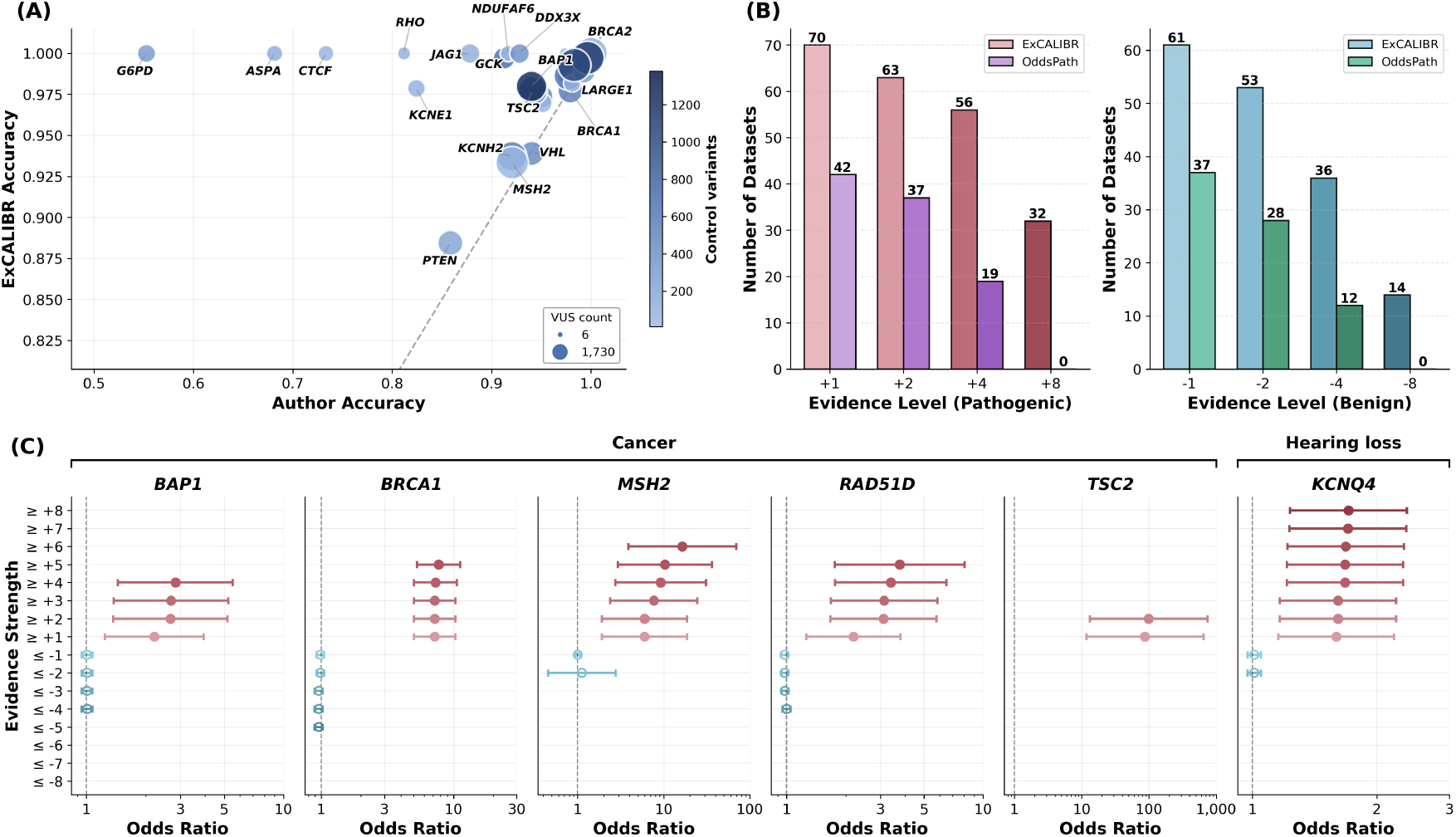
(A) Accuracy comparison of ExCALIBR out-of-bag evidence versus author annotations across the 28 genes with author-provided classifications. Two genes (*CRX* and *CARD11* ) were excluded from visualization as ExCALIBR assigned indeterminate evidence to all variants due to insufficient discriminatory power from their assays (20 and 11 control variants, respectively, with author accuracies of 0.750 and 0.857). The dashed line denotes equal accuracy. (B) Comparison of how many datasets reached each evidence strength for ExCALIBR vs. the approach described in Brnich *et al*. [12], often referred to as the OddsPath approach. Left: pathogenic evidence strengths; right: benign evidence strengths. (C) Odds ratios for occurrence of disease in individuals with variants meeting each ExCALIBR-calibrated evidence strength threshold in the All of Us biobank for six example genes. Genes are stratified by the type of disease association. Circles denote the estimated odds ratio and whiskers denote the 95% confidence interval. Filled circles denote estimates whose 95% confidence interval does not include 1; open circles denote those whose interval includes 1. The x-axes show odds ratios on a log scale.

Finally, we replicated the analysis from Figure 4 on the same set of 39 datasets with author-provided annotations using P/LP and B/LB variants from the ClinGen Evidence Repository [28]. To avoid circularity, we removed all experimental evidence (PS3/BS3 criteria) that contributed to classifications, resulting in 255 P/LP and 21 B/LB variants spanning 8 genes retaining enough evidence to remain P/LP or B/LB, respectively. ExCALIBR again outperformed author-provided annotations across all metrics, with no B/LB variants misclassified as pathogenic (Supplementary Figure S3 and Table S4). The 10 P/LP variants assigned benign evidence by ExCALIBR all originated from *PTEN* assays and received only weak benign strengths, with 7 at supporting (−1) and 3 at moderate (−2). These results demonstrate that ExCALIBR substantially improves accuracy over categorical threshold-based approaches and provides a basis for improved variant classification.

### ExCALIBR illuminates assay discriminatory power

The distribution of evidence strengths assigned by ExCALIBR across individual datasets provides insight into the discriminatory power of each experimental assay (Figure 6). Datasets whose variants are predominantly assigned indeterminate evidence demonstrate limited ability to distinguish functionally normal from abnormal variants. Conversely, datasets that reach strong pathogenic or benign evidence exemplify the assay’s ability to separate these two clinical classes.

**Figure 6.**
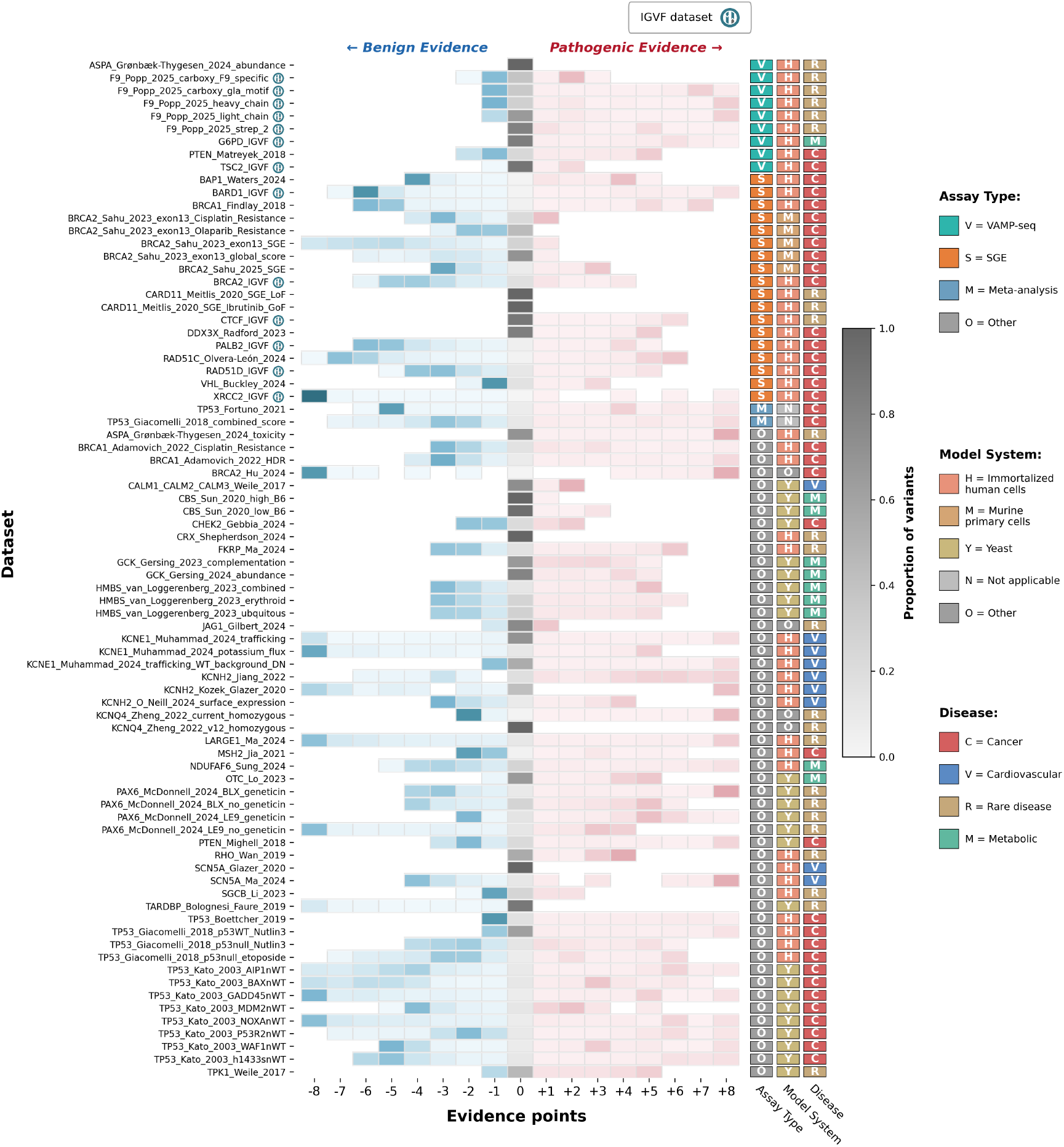
Distribution of ExCALIBR out-of-bag evidence strength assignments for each assay, grouped by the assay type. IGVF-produced datasets are labeled alongside the assay type, model system, and disease type. All possible evidence strengths between −8 and +8 are shown.

VAMP-seq assays tend to assign primarily pathogenic evidence, consistent with the fact that these assays measure protein abundance and do not consider all pathomechanisms [29]. For example, eight of nine VAMP-seq datasets reached a pathogenic evidence strength of moderate (+2), while only two reached benign evidence of the same strength; six reached strong pathogenic (+4), while none reached strong benign (−4). Across all VAMP-seq assays, 15.0% of variants reached pathogenic evidence of +2, compared to only 0.7% reaching benign evidence of −2. This asymmetry reflects the inherent limitation of measuring a single aspect of protein function while highlighting VAMP-seq’s strength in identifying damaging variants likely to be pathogenic.

Saturation Genome Editing (SGE) assays, by contrast, typically produce evidence strengths spanning both pathogenic and benign directions, reflecting their direct measurement of cellular fitness that covers a broader range of pathomechanisms [22]. Twelve of 18 SGE datasets reached pathogenic evidence of +2, and 14 reached benign evidence of −2. At the strong evidence level, 10 reached +4 and 11 reached −4. This illustrates the bidirectional discriminatory power of SGE.

We note that the observed discriminatory power also reflects the concordance between the control variants used for calibration and the disease mechanism measured by the assay. For genes associated with multiple diseases or phenotypes, ClinVar controls may be aggregated across phenotypes and not reflect the disease mechanism measured by the assay, artificially degrading calibration performance and thus the observed discriminatory power. Disentangling these effects at scale would require machine-readable, phenotype-specific annotations of control variants, which are not currently available.

Overall, our analysis demonstrates that the distribution of evidence strengths assigned by ExCALIBR can serve as a quantitative measure of assay discriminatory power and inform the utility of each assay for clinical variant classification, though interpretation should account for the match between control variants and assay mechanism.

## Discussion

High-throughput functional assays are a valuable tool for the clinical classification of variants [18]. To ensure consistent and accurate application of experimental evidence, we present a semi-supervised statistical framework and a method (ExCALIBR) for calibrating high-throughput functional assays. This framework jointly models assay score distributions of pathogenic, benign and synonymous variants along with that of a reference set of variants from gnomAD, as unique mixtures of skew normal distributions. The model computes the local positive likelihood ratio along with the prior probability of pathogenicity for variants in a gene or domain, all of which are essential components of calibration. These parameters are then directly translated into the quantitative evidence framework compatible with the ACMG/AMP clinical guidelines [7].

The results demonstrate the suitability of skew normal mixture models to faithfully capture assay score distributions from a diverse set of high-throughput functional experiments. ExCALIBR rigorously quantifies the strength of experimental evidence in clinical variant classification, and its distinct feature is that the evidence strength assigned to variants is variant-specific. Our model enables up to 16 distinct evidence strengths (±1, ±2, …, ±8) for the same assay, compared to at most two under current recommendations [12].

This calibration approach is limited by its dependence on the availability of an unbiased sample of known pathogenic and benign variants to accurately compute local positive likelihood ratios. The modeling does not account for potential biases in the collection of classified variants; e.g., the possibility of known pathogenic variants being more deleterious and having larger functional effects than pathogenic variants overall. Further work is needed to detect and quantify the severity of bias in assay distributions and correct for the effect of this bias on calibration [30–32]. While we limit circularity for genes where experimental assays considered in this work were already used to assert variant pathogenicity in ClinVar, experimental data from other assays may have been incorporated into classifying some of the pathogenic and benign variants, potentially resulting in a secondary circularity effect. Such detailed application of evidence is inconsistently recorded in ClinVar, making it difficult to unequivocally account for circularity. Thus, future calibration approaches must continue to be aware of potential risks of circularity. Additionally, while the skew normal distributions capture the distributions of the high-throughput experimental data analyzed here, the quality of the model fit may vary on some datasets and is affected by the normalization steps taken in generating experimental scores. Finally, the calibration approach has not been evaluated on genes with both gain- and loss-of-function disease mechanisms; our framework could be extended to handle such cases as relevant experimental datasets become available.

To our knowledge, this work presents the first approach to calibration *sensu stricto* for clinical interpretation of high-throughput functional assays. More broadly, calibration is a well-studied problem in machine learning, where the objective is to adjust the scores of trained classifiers to properly reflect posterior probabilities [33–35]. Typical algorithms in this space rely on relatively large quantities of data and, more importantly, assume the labeled training set is representative of the target distribution. This assumption does not hold for ClinVar controls, which are enriched for pathogenic variants through clinical ascertainment bias and do not reflect population-level frequencies of pathogenicity. Therefore, we introduced a reference set (gnomAD) to estimate the class prior in a population-aware manner, performing calibration in a semi-supervised framework.

Calibration has also been studied in the context of variant interpretation [36, 37]. Pejaver *et al*. [38] proposed a nonparametric method for the calibration of computational tools, but this model relies on larger quantities of control pathogenic and benign variants, which are unavailable for most genes. Therefore, stronger assumptions must be adopted to constrain the calibration. In terms of high-throughput experimental data, assay score distributions have previously been modeled with multi-sample Gaussian mixtures (MSGMM), learned in the positive-unlabeled classification setting to assign discrete functional annotations to variants [39]. Although not addressed in that work, the distributions learned by the MSGMM can be used to calculate a variant’s posterior probability of being functionally abnormal. However, functional abnormality does not equate to pathogenicity, and without restrictive assumptions on the functional mechanisms of disease, these probabilities alone are insufficient to calibrate assay scores to posterior probabilities of pathogenicity. Additional methods, such as MaveLLR [40] and acmgscaler [41], have been proposed to quantify the evidence strength of assay scores. These methods approximate the assay score distributions of pathogenic and benign variants using kernel density estimation (KDE), and these density functions are subsequently used to compute the local positive likelihood ratio at each assay score. While MaveLLR and acmgscaler can perform well in the presence of a large clinical control set, KDE is sensitive to the choice of kernel with limited data. These approaches also do not estimate the prior probability of pathogenicity, a value necessary for computing the posterior probability. These limitations prevent rigorous calibration and reliable evidence assignments.

Overall, our work demonstrates the feasibility of calibrating high-throughput functional assays for clinical variant classification and provides a method and software for this purpose. We find that the existing assays are sufficiently reliable to contribute to variant classification and thus reduce the number of rapidly accumulating VUS. The calibration of experimental assays will lead to an increased accuracy of genetic diagnosis and improved medical management for individuals affected by Mendelian disorders.

## Methods

The objective in calibrating high-throughput experimental data for a given gene or domain is to quantify the strength of evidence an assay score can serve in classifying a specific variant as pathogenic or benign. That is, we seek to learn a mapping function from a continuous assay score to a discrete evidence strength (supporting, moderate, strong, very strong) and direction (pathogenic, benign). Variants whose scores do not map to any evidence strength are said to have “indeterminate” strength and direction [38].

### Calibration framework

To learn the calibration function, we follow the general probabilistic framework based on the relationship between prior and posterior odds of pathogenicity [42]:

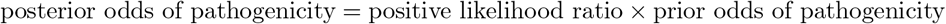

Letting *E* be a discrete random variable representing evidence and *Y* be a binary random variable representing whether (*Y* = 1) or not (*Y* = 0) a variant causes rare disease, the posterior odds of pathogenicity can be expressed as *P* (*Y* =1|*E*=*e*)*/P* (*Y* =0|*E*=*e*), the prior odds of pathogenicity as *P* (*Y* =1)*/P* (*Y* =0), and the positive likelihood ratio LR^+^(*e*) as *P* (*E*=*e*|*Y* =1)*/P* (*E*=*e*|*Y* =0).

To model the expert-derived rules from the ACMG/AMP guidelines [1], Tavtigian *et al*. [6] proposed to express LR^+^ in an exponential form as

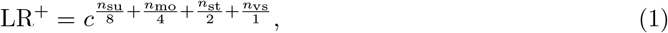

where *c* is a parameter and *n*_su_, *n*_mo_, *n*_st_, and *n*_vs_ are the numbers of supporting, moderate, strong, and very strong lines of evidence, respectively. Equation (1) specifies that one line of very strong evidence equals two, four, and eight lines of strong, moderate, and supporting evidence, respectively. This can be interpreted as a point-based system in which supporting, moderate, strong, and very strong evidence of pathogenicity or benignity are designated ± 1, ± 2, ± 4, ± 8 points, respectively, with positive points indicating pathogenic evidence and negative points denoting benign evidence [7].

The parameter *c* is determined as the smallest number in which the posterior probability of pathogenicity

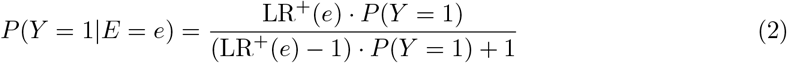

satisfies the classification thresholds from Richards *et al*. [1]: *P* (*Y* = 1|*E* = *e*) ≥ 0.9 for likely pathogenic and *P* (*Y* = 1|*E* = *e*) ≥ 0.99 for pathogenic variants. A prior of *P* (*Y* = 1) = 0.1 implies *c* = 350 [6], whereas *P* (*Y* = 1) = 0.044 implies *c* = 1124 [38].

When the available evidence is real-valued, it is necessary to discretize it for compatibility with ACMG/AMP guidelines [1]. Letting *S* be a real-valued random variable representing a variant’s assay score, we estimate the local positive likelihood ratio lr^+^(*s*) at score *s* as the density ratio *p*(*s*|*Y* =1)*/p*(*s*|*Y* =0) and establish score intervals [τ_low_, τ_high_] such that lr^+^(*s*) is greater than the required level for all scores in the interval [38]. For example, supporting evidence requires 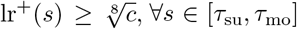, *∀s ∈* [τ_su_, τ_mo_], while moderate evidence requires 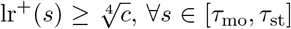, *∀s ∈* [τ_mo_, τ_st_]. Finer-grained levels of evidence (e.g., ± 3, ± 5 points) can be modeled in the same manner.

To calibrate *in silico* tools, Pejaver *et al*. [38] combined the data from all disease-associated genes and used separate nonparametric methods to estimate *P* (*Y* = 1) and lr^+^(*s*). However, this method relies on a relatively large number of clinical control variants and cannot be reliably applied to most individual genes. We therefore developed a flexible parametric approach to simultaneously estimate *P* (*Y* = 1) and lr^+^(*s*) in a gene-specific manner.

### Data

As part of the IGVF Consortium, a harmonized set of 83 experimental datasets across 40 clinically relevant genes was recently developed according to Tejura *et al*. [21]. This harmonized resource contains variant effect measurements for 255,354 unique variants, consisting of newly generated data through IGVF, curated experimental datasets from MaveDB [16] and published literature, and meta-analyses with classifier models aggregating scores across different experiments for a given gene.

A typical dataset contains continuous scores measuring the functional effect of hundreds to thousands of variants, including single and multiple nucleotide substitutions, insertions, and deletions. These scores may quantify variant effects on specific protein properties (abundance, cell surface localization) or overall cell fitness. Additionally, 19 assays used endogenous genome editing and therefore detect splice-altering variants, whereas 64 relied on cDNA-based approaches that assess functional consequences solely at the amino acid level.

In this study, we calibrated 80 datasets across 39 genes (Supplementary Data 1), excluding two meta-analyses containing binary classifier outputs for *F9* [43] and *TP53* [44] variants, respectively, and a newly generated dataset of *SFPQ* variants [21] given the absence of a validated gene-disease association. Some experimental datasets include score intervals that specify functional annotations for each variant (broadly categorized as functionally normal, functionally abnormal, or indeterminate), which we used for comparison with our method. These intervals can be determined using arbitrary thresholds, predictive models, or a combination of approaches. In addition, four datasets were post-processed to define score intervals as in Woo *et al*. [45] (Supplementary Data 1).

Clinical controls were collected from ClinVar [2] (January 2025). Variant subsets, referred to as P/LP, B/LB, and VUS, were constructed by selecting variants with review status of at least one star and clinical significance of pathogenic (P) or likely pathogenic (LP), benign (B) or likely benign (LB), and uncertain significance (VUS), respectively. Population variants in gnomAD v4.1.0 genomes and exomes [19] were also collected. For assays that do not detect splice-altering effects of variants, any variant with a SpliceAI [46] score, using the maximum of the acceptor loss/gain and donor loss/gain predictions, greater than 0.2 [47] was removed, as it is unknown if any associated pathogenic effect is mediated by disrupted protein function or aberrant splicing.

### Modeling

Assay score distributions of P/LP, B/LB, and gnomAD variants from each experimental dataset are approximated using a multi-sample skew normal mixture model; for details refer to Supplementary Section 1.1. Skew normal distributions are chosen as they are able to model the asymmetry observed in experimental datasets and have been found to be tractably incorporated into multi-sample mixture models [48]. We model each score distribution with both a two-component and three-component mixture. In the former case, these components represent functionally abnormal and normal variant assay scores. In some datasets, the third component is incorporated to represent a hypomorphic region or for model flexibility; see Supplementary Section 1.2 for parameterization details. All parameters are jointly learned by adapting the expectation-maximization algorithm derived by Peng *et al*. [48]. We additionally enforce a relaxed density constraint between components to promote monotonicity of the positive likelihood ratio. Optimization details are described in Supplementary Section 1.3.

At each assay score *s*, we estimate the local positive likelihood ratio lr^+^(*s*) as the ratio of pathogenic to benign sample mixture densities at *s* (Supplementary Section 1.4). Next, we estimate the prior probability of pathogenicity *P* (*Y* = 1) using an expectation-maximization algorithm adapted from methods for label shift correction [49] that iteratively computes posterior probabilities for population variants and averages them (Supplementary Section 1.5). Finally, if available, synonymous variant scores are also modeled to aid in parameter optimization and to represent a functionally normal distribution, under the assumption that synonymous variants are predominantly functionally normal. Synonymous variant scores were available for 53 of 80 datasets.

Samples are constructed to be mutually exclusive for model fitting. Variants eligible for multiple samples are assigned at random to one, with the exception that synonymous variants are preferentially assigned to the synonymous sample when available.

### Evidence assignment

Uncertainty is quantified through 1,000 bootstrap iterations of the dataset, with each sample (P/LP, B/LB, gnomAD, and synonymous) resampled independently. In this work, we mainly present evidence assigned in an out-of-bag manner; however, ExCALIBR uses all 1,000 bootstrap iterations by default. To assign robust evidence throughout the score range, the 5^th^ and 95^th^ percentiles of lr^+^(*s*), along with the median valid prior, are used to determine pathogenic and benign evidence strengths, respectively. Subsequent refinements ensure monotonicity of evidence assignments throughout the score range if not already satisfied. We assign evidence points along all integers between −8 and +8, providing more granular evidence assignment than current recommendations [12].

### Model selection

Each dataset can be modeled using a choice of two-component versus three-component mixtures, relaxed versus strict density constraint, evidence assignment refinement strategies, and various options for the benign reference (necessary for prior and lr^+^ estimation): benign sample only, synonymous sample only, or averaged benign and synonymous sample mixing proportions. In this study, we trained all model configurations and selected the final model (Supplementary Data 1) based on both automated selection criteria and manual evaluation of fit quality and evidence assignments. Selection criteria are described in Supplementary Section 1.6.

### Missing samples

In the case of a dataset lacking a positive (pathogenic) or negative (benign and synonymous) sample, but not both, we adapt our methods for computing the lr^+^ and prior. When neither benign nor synonymous variants are available, we use a positive-unlabeled (PU) learning framework; when benign or synonymous variants are available but pathogenic variants are unavailable, we use a negative-unlabeled (NU) learning framework; see Supplementary Section 1.7 for details. In the case of a missing benign sample but not synonymous, synonymous variants serve as the benign control. The population sample (here gnomAD, but user-defined in general) is always needed because the prior probability of pathogenicity is essential for calibration.

### Evaluation

#### Model fit quality

We quantified model fit quality using the normalized Yang *et al*. [27] distance (*p* = 2) between the empirical and fitted cumulative distribution functions for each sample. We use this distance metric rather than likelihood, as likelihood values are not as interpretable or comparable across datasets. The distance ranges from 0 to 1 and is calculated using real-valued vectors at all unique scores from the bootstrapped assay.

### Biobank validation

We assessed the relationship between ExCALIBR evidence strengths and clinical phenotypes using approximately 400,000 participants from the All of Us biobank. Validation was performed using the All of Us Workbench and the Controlled Tier Dataset v8 [15].

Of the 39 clinically-relevant genes in this study, we identified 17 autosomal dominant genes with sufficient biobank data for validation: those associated with phenotypes definable through All of Us diagnosis codes, measurements, or surveys, and with at least 40 whole-genome sequenced participants having variants in the gene. The remaining gene-disease pairs were excluded due to non-autosomal-dominant inheritance, absence of a matching phenotype in All of Us, or insufficient participant counts (Supplementary Data 4). For each included gene, we defined case cohorts as participants that had specific conditions, measurements, or survey answers that indicated they had the phenotype of interest, and control cohorts by excluding participants that had broad conditions, measurements, or survey answers that indicated they might have the phenotype of interest or a closely related phenotype (Supplementary Data 4).

We grouped variants by ExCALIBR-assigned evidence strength and queried the All of Us short-read whole genome sequencing matrix to identify carriers, defined as participants with one or more variants in each set. For each evidence strength, we estimated the odds ratio of disease given carrier status by fitting logistic regression models with case/control status as the response variable. Explanatory variables included carrier status, sex assigned at birth (male, female, or other), age at data release, and 16 genetic ancestry principal components computed by All of Us [50]. For assays that did not detect splice-altering effects, we excluded participants with variants having SpliceAI scores above 0.2 using the maximum of acceptor loss/gain and donor loss/gain predictions. We report point estimates, 95% confidence intervals, and Wald test p-values for the regression coefficient of the carrier status variable, which represents the natural log of the odds ratio (Supplementary Data 4).

## Supporting information

Dataset Summaries

Model Fit Visualizations

OddsPath Evidence

All of Us Biobank Validation

## Acknowledgements

We thank the ClinGen Sequence Variant Interpretation Working Group for their guidance. This work has been supported by the NIH awards U01 HG012022 (PR), R00 LM012992 (VP), U24 HG011450 (AODL), U01 HG011755 (AODL), U01 HG012039 (MC), R35 GM152106 (DMF), and UM1 HG011969 (DMF, LMS); an Early Career Award from Alex’s Lemonade Stand for Childhood Cancer and the RUNX1 Foundation 21-25037 (AEM), and the Brotman Baty Institute Catalytic Collaborations Grant CC28 (AEM). This work was completed in part using the Explorer Cluster, supported by Northeastern University’s Research Computing team.

## Competing Interests

The authors declare the following competing interests. D.M.F. is a member of the Alloz Bio scientific advisory board. The remaining authors declare no competing interests.

## Code and Data Availability

Source code is available at https://github.com/rosstewart/exCALIBR. Datasets used in this study are further described in Tejura *et al*. [21]. Assay score distributions and visualized evidence thresholds for each dataset are available in MaveDB [16] and MaveMD [51]. Evidence thresholds (IGVF accession no. IGVFFI5038ZNHZ), variant-level evidence assignments per assay (IGVF accession no. IGVFFI7610PCPU), and per-variant aggregated experimental and predictive evidence (IGVF accession no. IGVFFI1443TQDN; predictive evidence derived from calibration of computational predictors in Chen *et al*. [52]) resulting from this work are available on the IGVF data portal [53].

## Notes

### Summary of Updates

This version of the manuscript includes additional methods and results.

