## Supplementary Information for "Gene-based calibration of high-throughput functional assays for clinical variant classification"

[1]Khoury College of Computer Sciences, Northeastern University, Boston, MA 02115, USA; [2]Department of Genome Sciences, University of Washington, Seattle, WA 98195, USA; [3]Brotman Baty Institute for Precision Medicine, University of Washington, Seattle, WA 98195, USA; [4]Department of Laboratory Medicine and Pathology, University of Washington, Seattle, WA 98195, USA; [5]Department of Biostatistics and Medical Informatics, University of Wisconsin, Madison, WI 53726, USA; [6]Institute for Genomic Health and Department of Genetics and Genomic Sciences, Icahn School of Medicine at Mount Sinai, New York, NY 10029, USA; [7]Bioinformatics Division, The Walter and Eliza Hall Institute of Medical Research, Parkville, VIC, Australia; Department of Medical Biology, University of Melbourne, Melbourne, VIC, Australia; [8]Department of Bioengineering, University of Washington, Seattle, WA 98195, USA; [9]Program in Medical and Population Genetics, Broad Institute of MIT and Harvard, Cambridge, MA 02142, USA; [10]Division of Genetics and Genomics, Boston Children's Hospital, Harvard Medical School, Boston, MA 02115, USA.

[†]These authors contributed equally to this work.

**2 Proof of Monotonicity of the Local Positive Likelihood Ratio** **5**

**3 Supplementary Figures** **6**

### 1 Modeling details

#### 1.1 Skew normal distributions

Assay scores are modeled using skew normal distributions. Under the distribution's canonical parameterization [1], the probability density function of a random variable $X \sim \text{SN}(\mu, \omega, \lambda)$ is given by

$$\mathrm{SN}(x;\mu,\omega,\lambda) = \frac{2}{\omega}\phi\left(\frac{x-\mu}{\omega}\right)\Phi\left(\lambda\left(\frac{x-\mu}{\omega}\right)\right), \tag{S1}$$

where $\mu$, $\omega$, and $\lambda$ denote the location, scale and skew of the distribution, and $\phi$ and $\Phi$ are the probability density and cumulative distribution functions of the standard Gaussian distribution, respectively. Under the alternate parameterization, defined by parameters $\Delta$ and $\Gamma$, used in the parameter updates of the optimization algorithm, $X \sim \mathrm{SN}(\mu,\omega,\lambda)$ can be expressed as $X = \mu + \Delta \cdot T + \Gamma^{\frac{1}{2}} \cdot U$, where $T$ is a random variable from the standard normal distribution truncated below 0 and U is a random variable from the standard normal distribution [2, 3]. Canonical and alternate parameterizations are related through equations in Table 1 of Peng *et al.* [3], and are included in Supplementary Table S5 for completeness.

#### 1.2 Parameterization

Let $S_P$ be the set of assay scores for variants in the dataset's pathogenic and likely pathogenic (P/LP) sample, let $S_B$ be the set of assay scores for variants in the dataset's benign and likely benign (B/LB) sample, and let $S_G$ be the set of assay scores for variants in the dataset's gnomAD sample, used here as a population reference.

Functionally abnormal ($a$), functionally normal ($n$), and hypomorphic ($h$) variant assay scores are parameterized by $\theta_a = (\mu^{(a)}, \omega^{(a)}, \lambda^{(a)})$, $\theta_n = (\mu^{(n)}, \omega^{(n)}, \lambda^{(n)})$, and $\theta_h = (\mu^{(h)}, \omega^{(h)}, \lambda^{(h)})$, respectively. The mixture model approximating the distributions of $S_P$, $S_B$, and $S_G$ can therefore be expressed as

$$p_{S_i}(s;\alpha_{S_i},\beta_{S_i},\theta_a,\theta_h,\theta_n) = \alpha_{S_i}\cdot\mathrm{SN}(s;\theta_a) + \beta_{S_i}\cdot\mathrm{SN}(s;\theta_h) + (1-\alpha_{S_i}-\beta_{S_i})\cdot\mathrm{SN}(s;\theta_n), \tag{S2}$$

where $\alpha_{S_i}$ and $\beta_{S_i}$ are the mixture weights (proportions) of the functionally abnormal and hypomorphic components, respectively, for sample $i \in \{P, B, G\}$, with $\alpha_{S_i}, \beta_{S_i} \in [0, 1]$ and $\alpha_{S_i} + \beta_{S_i} \leq 1$. In the case of a two-component model, $\beta_{S_i}$ is simply zero. If available, the sample of synonymous variant assay scores $S_S$ is similarly represented with mixture weights $(\alpha_{S_S}, \beta_{S_S})$.

#### 1.3 Optimization

##### 1.3.1 Parameter update equations

With $\ddot{}$ and $\bar{}$ denoting new and old parameter values, respectively, the parameter update equations can be expressed as

$$\ddot{\mu}^{(*)} = \frac{\sum_{S_i\in\{S_P,S_B,S_G,S_S\}}\sum_{s_{ij}\in S_i}\bar{p}^{(*)}_{S_i}(s_{ij})\cdot\bar{m}^{(*)}\left(s_{ij},\bar{\Delta}^{(*)}\right)}{\sum_{S_i\in\{S_P,S_B,S_G,S_S\}}\sum_{s_{ij}\in S_i}\bar{p}^{(*)}_{S_i}(s_{ij})}$$

$$\ddot{\Delta}^{(*)} = \frac{\sum_{S_i\in\{S_P,S_B,S_G,S_S\}}\sum_{s_{ij}\in S_i}\bar{p}^{(*)}_{S_i}(s_{ij})\bar{d}^{(*)}(s_{ij},\ddot{\mu}^{(*)})}{\sum_{S_i\in\{S_P,S_B,S_G,S_S\}}\sum_{s_{ij}\in S_i}\bar{p}^{(*)}_{S_i}(s_{ij})}$$

$$\ddot{\Gamma}^{(*)} = \frac{\sum_{S_i\in\{S_P,S_B,S_G,S_S\}}\sum_{s_{ij}\in S_i}\bar{p}^{(*)}_{S_i}(s_{ij})\bar{g}^{(*)}(s_{ij},\ddot{\mu}^{(*)},\ddot{\Delta}^{(*)})}{\sum_{S_i\in\{S_P,S_B,S_G,S_S\}}\sum_{s_{ij}\in S_i}\bar{p}^{(*)}_{S_i}(s_{ij})}$$

with $* \in \{a, h, n\}$. $\bar{p}_{S_i}^{(a)}$, $\bar{p}_{S_i}^{(h)}$, and $\bar{p}_{S_i}^{(n)}$ are the component posterior probabilities (responsibilities) defined as:

$$\bar{p}_{S_i}^{(a)}(s) = \frac{\bar{\alpha}_{S_i} \cdot \mathrm{SN}(s; \bar{\theta}_a)}{\bar{\alpha}_{S_i} \cdot \mathrm{SN}(s; \bar{\theta}_a) + \bar{\beta}_{S_i} \cdot \mathrm{SN}(s; \bar{\theta}_h) + (1 - \bar{\alpha}_{S_i} - \bar{\beta}_{S_i}) \cdot \mathrm{SN}(s; \bar{\theta}_n)}$$

$$\bar{p}_{S_i}^{(h)}(s) = \frac{\bar{\beta}_{S_i} \cdot \mathrm{SN}(s; \bar{\theta}_h)}{\bar{\alpha}_{S_i} \cdot \mathrm{SN}(s; \bar{\theta}_a) + \bar{\beta}_{S_i} \cdot \mathrm{SN}(s; \bar{\theta}_h) + (1 - \bar{\alpha}_{S_i} - \bar{\beta}_{S_i}) \cdot \mathrm{SN}(s; \bar{\theta}_n)}$$

$$\bar{p}_{S_i}^{(n)}(s) = \frac{(1 - \bar{\alpha}_{S_i} - \bar{\beta}_{S_i}) \cdot \mathrm{SN}(s; \bar{\theta}_n)}{\bar{\alpha}_{S_i} \cdot \mathrm{SN}(s; \bar{\theta}_a) + \bar{\beta}_{S_i} \cdot \mathrm{SN}(s; \bar{\theta}_h) + (1 - \bar{\alpha}_{S_i} - \bar{\beta}_{S_i}) \cdot \mathrm{SN}(s; \bar{\theta}_n)}$$

The weight updates are computed as:

$$\ddot{\alpha}_{S_i} = \frac{1}{|S_i|} \sum_{s_{ij} \in S_i} \bar{p}_{S_i}^{(a)}(s_{ij}), \quad \ddot{\beta}_{S_i} = \frac{1}{|S_i|} \sum_{s_{ij} \in S_i} \bar{p}_{S_i}^{(h)}(s_{ij}), \quad \forall i \in \{P, B, G, S\} \tag{S3}$$

where the weights represent the mean posterior probability (responsibility) of each component across all observations in sample $i$. $\bar{m}^{(*)}(s, \Delta)$, $\bar{d}^{(*)}(s, \mu)$, and $\bar{g}^{(*)}(s, \mu, \Delta)$ are defined in Supplementary Table S6.

###### 1.3.2 Density constraint

We enforce a relaxed monotonicity constraint on the density ratios between consecutive components:

$$\frac{\mathrm{SN}(s_1; \theta_a)}{\mathrm{SN}(s_1; \theta_h)} \geq \frac{\mathrm{SN}(s_2; \theta_a)}{\mathrm{SN}(s_2; \theta_h)}, \quad \forall s_1 \leq s_2 \tag{S4}$$

$$\frac{\mathrm{SN}(s_1; \theta_h)}{\mathrm{SN}(s_1; \theta_n)} \geq \frac{\mathrm{SN}(s_2; \theta_h)}{\mathrm{SN}(s_2; \theta_n)}, \quad \forall s_1 \leq s_2 \tag{S5}$$

These constraints are only enforced in regions where both component densities are non-negligible ($> e^{-7}$), allowing flexibility in the distribution tails. This pairwise monotonicity between consecutive components, combined with the assumption that $\alpha_{S_P} > \alpha_{S_B}$ (pathogenic variants are enriched for functionally abnormal variants relative to benign variants, as verified during assay validation [4]), promotes monotonicity of the positive likelihood ratio $\mathrm{lr}^+(s)$. In the case of an assay that scores functionally abnormal variants higher than functionally normal variants, these parameterizations can simply be considered flipped.

**Theorem 1.** *For a two-component model ($\beta_{S_i} = 0$), given a monotonic density ratio $SN(s;\theta_a)/SN(s;\theta_n)$ in regions of non-negligible density and $\alpha_{S_P} > \alpha_{S_B}$, the likelihood ratio $p_{S_P}(s)/p_{S_B}(s)$ is monotonic in those regions.*

The proof is in Supplementary Section 2. While these constraints promote monotonicity of the positive likelihood ratio $\mathrm{lr}^+(s)$, violations may occur with a three-component model and in low-density regions where the constraint is not enforced. As done by Peng *et al.* [5], a binary search is used to maintain these constraints at each update to the component parameters $\mu^{(*)}, \Delta^{(*)}, \Gamma^{(*)}$ for $* \in \{a, h, n\}$.

###### 1.3.3 Initialization

To initialize the component parameters $\theta_a$, $\theta_h$, and $\theta_n$, we use two complementary approaches: (1) K-means clustering with $K \in \{2, 3\}$ (matching the model choice) applied to all observations, or

(2) method of moments estimation on random partitions of the data. For K-means initialization, normal distributions are fit to each cluster to obtain initial location and scale parameters $\mu^{(*)}$ and $\omega^{(*)}$. For method of moments initialization, observations are randomly partitioned into $K$ groups and moments are computed for each partition. As it has been observed empirically that the direction of the skew does not change during the iterations of the EM algorithm [3], skewness parameters $\lambda^{(*)}$ are randomly sampled from $\mathbb{U}(-0.25, 0.25)$ for K-means initialization. For method of moments, they are estimated from sample skewness with fallback to uniform sampling if estimation fails.

To ensure initial parameters satisfy the density constraints (Eqs. S4–S5), component scale parameters $\omega^{(*)}$ are iteratively reduced until all constraints are satisfied within the observed score range. To mitigate the effects of random initialization, we perform 100 independent runs of the modeling procedure, with each run randomly selecting one of the two initialization strategies. The model that yields the highest likelihood across all runs is retained.

#### 1.4 Local positive likelihood ratio

$$\mathrm{lr}^{+}(s; \alpha_{S_P}, \beta_{S_P}, \alpha_{S_B}, \beta_{S_B}, \theta_a, \theta_h, \theta_n) = \frac{p_{S_P}(s; \alpha_{S_P}, \beta_{S_P}, \theta_a, \theta_h, \theta_n)}{p_{S_B}(s; \alpha_{S_B}, \beta_{S_B}, \theta_a, \theta_h, \theta_n)} \tag{S6}$$

#### 1.5 Prior estimation

The prior probability of pathogenicity $P(Y = 1)$ is estimated using an expectation-maximization algorithm adapted from methods for label shift correction [6]. Given the fitted mixture model densities $p_{S_P}(s)$ and $p_{S_B}(s)$ for pathogenic and benign score distributions, we initialize $P(Y = 1) = 0.5$ and iteratively update via:

$$P(Y = 1)^{(t+1)} = \frac{1}{|S_G|} \sum_{s \in S_G} \frac{1}{1 + \frac{1-P(Y=1)^{(t)}}{P(Y=1)^{(t)}} \cdot \frac{p_{S_B}(s)}{p_{S_P}(s)}}, \tag{S7}$$

where the sum is over all variants in the gnomAD reference set $S_G$, and the algorithm is run until convergence. Although we used gnomAD as a representative population set, ExCALIBR enables users to select an alternative reference set, such as all single-nucleotide missense variants. This may improve the stability of the modeling but could yield less accurate prior estimates. Users can also manually specify a prior when external data suggest a different value is appropriate.

#### 1.6 Model selection criteria

**Component selection.** In addition to manual fit and evidence inspection, we compare the validation log-likelihoods of each model trained on the same bootstrapped sample. The three-component model is preferred if at least 95% of bootstrap iterations show improvement, ensuring robust evidence of improved fit despite increased model capacity. Otherwise, the simpler two-component model is favored.

**Density constraint relaxation.** We typically prefer to relax the density constraint; i.e., allowing constraint violations in negligible density regions, which aids in modeling flexibility. However, we chose to strictly enforce this density constraint throughout the score range for 15 of 80 datasets where this yielded more reliable results (Supplementary Data 1).

**Evidence strength threshold post-processing.** When evidence assignments are not monotonic, we post-process the thresholds to ensure they are either consistently increasing or decreasing. This lack of monotonicity can occur in three-component models and in low-density regions due to our imposed optimization constraints. To enforce monotonicity, we extend evidence thresholds to the score range extremes when the defined evidence falls back to indeterminate. However, for three of 80 datasets (`DDX3X_Radford_2023`, `F9_Popp_2025_heavy_chain`, `GCK_Gersing_2023_complementation`) where this extension would assign evidence in unreliable regions, we instead remove the evidence threshold entirely.

**Benign reference selection.** We select the configuration (benign, synonymous, or average) that assigns the most appropriate or conservative evidence. When benign samples are limited, averaging benign and synonymous mixing proportions typically yields more reliable evidence.

##### 1.7 Positive-unlabeled and negative-unlabeled learning frameworks

In PU and NU learning scenarios, the gnomAD reference population $S_G$ is modeled as a mixture of the available labeled sample (pathogenic or benign) and the missing distribution (benign or pathogenic, respectively). Given that $p_{S_G}(s) = P(Y=1) \cdot p_{S_P}(s) + P(Y=0) \cdot p_{S_B}(s)$, we can recover the missing density distribution by rearrangement. For PU learning with only pathogenic labels available:

$$p_{S_B}(s) = \frac{p_{S_G}(s) - P(Y=1) \cdot p_{S_P}(s)}{1 - P(Y=1)}, \tag{S8}$$

and similarly for NU learning with only benign labels:

$$p_{S_P}(s) = \frac{p_{S_G}(s) - P(Y=0) \cdot p_{S_B}(s)}{P(Y=1)}. \tag{S9}$$

The prior probability is estimated using a modified EM algorithm. For PU learning, we initialize $P(Y=1) = 0.1$ and update via:

$$P(Y=1)^{(t+1)} = \frac{1}{|S_G|} \sum_{s \in S_G} \frac{P(Y=1)^{(t)} \cdot p_{S_P}(s)}{p_{S_G}(s)}. \tag{S10}$$

For NU learning, we estimate $P(Y=0)$ analogously (initialized at 0.9) and compute $P(Y=1) = 1 - P(Y=0)$. The likelihood ratio is then computed using the inferred density from Eqs. S8 or S9.

#### 2 Proof of Monotonicity of the Local Positive Likelihood Ratio

To prove Theorem 1, the local positive likelihood ratio is expressed as

$$\begin{aligned}
\mathrm{lr}^+(s; w_{S_P}, w_{S_B}, \theta_a, \theta_n) &= \frac{p_{S_P}(s; w_{S_P}, \theta_a, \theta_n)}{p_{S_B}(s; w_{S_B}, \theta_a, \theta_n)} \\
&= \frac{w_{S_P}\mathrm{SN}(s; \theta_a) + (1 - w_{S_P})\mathrm{SN}(s; \theta_n)}{w_{S_B}\mathrm{SN}(s; \theta_a) + (1 - w_{S_B})\mathrm{SN}(s; \theta_n)}
\end{aligned}$$

The derivative of $\mathrm{lr}^+(s; w_{S_P}, w_{S_B}, \theta_a, \theta_n)$ is computed as

$$\begin{aligned}
\frac{\mathrm{d}}{\mathrm{d}s}\mathrm{lr}^+(s; w_{S_P}, w_{S_B}, \theta_a, \theta_n) =& \frac{p_{S_B}(s; w_{S_B}, \theta_a, \theta_n)\left(w_{S_P}\mathrm{SN}'(s; \theta_a) + (1 - w_{S_P})\mathrm{SN}'(s; \theta_n)\right)}{p^2_{S_B}(s; w_{S_B}, \theta_a, \theta_n)} - \\
& \frac{p_{S_P}(s; w_{S_P}, \theta_a, \theta_n)\left(w_{S_B}\mathrm{SN}'(s; \theta_a) + (1 - w_{S_B})\mathrm{SN}'(s; \theta_n)\right)}{p^2_{S_B}(s; w_{S_B}, \theta_a, \theta_n)}
\end{aligned}$$

Combining and simplifying terms, this can be expressed as

$$\frac{\mathrm{d}}{\mathrm{d}s}\mathrm{lr}^+(s;w_{S_P},w_{S_B},\theta_a,\theta_n)=\frac{(w_{S_P}-w_{S_B})\Big(\mathrm{SN}(s;\theta_n)\mathrm{SN}'(s;\theta_a)-\mathrm{SN}(s;\theta_a)\mathrm{SN}'(s;\theta_n)\Big)}{p^2_{S_B;}(s;w_{S_B},\theta_a,\theta_n)}$$

As it is given that $\frac{\mathrm{SN}(s;\theta_a)}{\mathrm{SN}(s;\theta_n)}$ is monotonic, the derivative can be expressed as

$$\frac{\mathrm{d}}{\mathrm{d}s}\frac{\mathrm{SN}(s;\theta_a)}{\mathrm{SN}(s;\theta_n)}=\frac{\mathrm{SN}(s;\theta_n)\mathrm{SN}'(s;\theta_a)-\mathrm{SN}(s;\theta_a)\mathrm{SN}'(s;\theta_n)}{\mathrm{SN}^2(s;\theta_n)}$$

and this value is positive for all scores $s$ if $\frac{\mathrm{SN}(s;\theta_a)}{\mathrm{SN}(s;\theta_n)}$ is monotonically increasing and negative if monotonically decreasing. The derivative of the local positive likelihood ratio can be expressed as

$$\frac{\mathrm{d}}{\mathrm{d}s}\mathrm{lr}^+(s;w_{S_P},w_{S_B},\theta_a,\theta_n)=c(s)\cdot\frac{\mathrm{d}}{\mathrm{d}s}\frac{\mathrm{SN}(s;\theta_a)}{\mathrm{SN}(s;\theta_n)}$$

where $c(s)=\frac{(w_{S_P}-w_{S_B})\mathrm{SN}^2(s;\theta_n)}{p^2_{S_B}(s;w_{S_B},\theta_a,\theta_n)}>0\ \forall s$, given $w_{S_P}>w_{S_B}$, from which it follows that the local positive likelihood ratio is monotonic. To show the posterior is monotonic, the posterior can be expressed using Bayes' Rule as

$$\begin{aligned}P(Y=1|s)&=\frac{p(s|Y=1)P(Y=1)}{p(s|Y=1)P(Y=1)+p(s|Y=0)(1-P(Y=1))}\\&=\frac{1}{1+\frac{1-P(Y=1)}{P(Y=1)}\frac{p(s|Y=0)}{p(s|Y=1)}}\\&=\frac{1}{1+\frac{1-P(Y=1)}{P(Y=1)}\mathrm{lr}^+(s;w_{S_P},w_{S_B},\theta_a,\theta_n)^{-1}}\end{aligned}$$

The derivative can be expressed as

$$\begin{aligned}\frac{\mathrm{d}}{\mathrm{d}s}P(Y=1|s)&=\frac{\frac{1-P(Y=1)}{P(Y=1)}\mathrm{lr}^+(s;w_{S_P},w_{S_B},\theta_a,\theta_n)^{-2}\frac{\mathrm{d}}{\mathrm{d}s}lr^+(s;w_{S_P},w_{S_B},\theta_a,\theta_n)}{\Big(1+\frac{1-P(Y=1)}{P(Y=1)}\mathrm{lr}^+(s;w_{S_P},w_{S_B},\theta_a,\theta_n)^{-1}\Big)^2}\\&=c_2(s)\frac{\mathrm{d}}{\mathrm{d}s}lr^+(s;w_{S_P},w_{S_B},\theta_a,\theta_n)\end{aligned}$$

with $c_2(s)=\frac{\frac{1-P(Y=1)}{P(Y=1)}}{\mathrm{lr}^+(s;w_{S_P},w_{S_B},\theta_a,\theta_n)^2\Big(1+\frac{1-P(Y=1)}{P(Y=1)}\mathrm{lr}^+(s;w_{S_P},w_{S_B},\theta_a,\theta_n)^{-1}\Big)^2}>0\ \forall s$, showing the posterior is monotonic.

### 3 Supplementary Figures

| Author Category | Abnormal | | | | Indeterminate | | | | Normal | | | |
|---|---|---|---|---|---|---|---|---|---|---|---|---|
| Gene | $n$ | Path. | Ind. | Ben. | $n$ | Path. | Ind. | Ben. | $n$ | Path. | Ind. | Ben. |
| *ASPA* | 1,077 | 33.7 | 66.3 | 0 | 1,242 | 6.8 | 93.2 | 0 | 1,425 | 7.2 | 92.8 | 0 |
| *BAP1* | 4,759 | 82.7 | 4.2 | 13.0 | 444 | 0 | 0 | 100 | 8,962 | 0 | 0.1 | 99.9 |
| *BARD1* | 1,129 | 96.0 | 4.0 | 0 | 174 | 0 | 72.4 | 27.6 | 7,548 | 0 | 0 | 100 |
| *BRCA1* | 1,105 | 95.7 | 4.3 | 0 | 441 | 17.7 | 49.0 | 33.3 | 3,693 | 0 | 0.2 | 99.8 |
| *BRCA2* | 130 | 100 | 0 | 0 | 12 | 0 | 25.0 | 75.0 | 302 | 0 | 0 | 100 |
| *CARD11* | 1,239 | 0 | 100 | 0 | 1,555 | 0 | 100 | 0 | 1,050 | 0 | 100 | 0 |
| *CRX* | 365 | 0 | 100 | 0 | 168 | 0 | 100 | 0 | 1,056 | 0 | 100 | 0 |
| *CTCF* | 590 | 91.4 | 8.6 | 0 | 108 | 2.8 | 97.2 | 0 | 4,092 | 0.1 | 99.9 | 0 |
| *DDX3X* | 1,638 | 87.5 | 12.5 | 0 | 710 | 0 | 100 | 0 | 6,732 | 0.5 | 99.5 | 0 |
| *FKRP* | 1,474 | 81.0 | 19.0 | 0 | 0 | 0 | 0 | 0 | 2,968 | 0 | 13.0 | 87.0 |
| *G6PD* | 602 | 90.5 | 9.5 | 0 | 94 | 0 | 100 | 0 | 2,763 | 0 | 100 | 0 |
| *GCK* | 2,162 | 72.9 | 27.1 | 0 | 2,851 | 0 | 100 | 0 | 849 | 0 | 100 | 0 |
| *JAG1* | 483 | 83.4 | 16.6 | 0 | 0 | 0 | 0 | 0 | 2,307 | 1.7 | 88.6 | 9.6 |
| *KCNE1* | 822 | 44.4 | 54.5 | 1.1 | 579 | 0 | 20.6 | 79.4 | 1,159 | 0 | 51.2 | 48.8 |
| *KCNH2* | 1,998 | 84.2 | 15.8 | 0 | 109 | 0 | 0 | 100 | 5,388 | 0 | 6.9 | 93.1 |
| *KCNQ4* | 916 | 65.0 | 30.3 | 4.7 | 284 | 100 | 0 | 0 | 6,108 | 0 | 50.1 | 49.9 |
| *LARGE1* | 1,380 | 100 | 0 | 0 | 0 | 0 | 0 | 0 | 5,323 | 1.3 | 14.7 | 84.0 |
| *MSH2* | 382 | 72.0 | 28.0 | 0 | 0 | 0 | 0 | 0 | 4,648 | 0 | 1.0 | 99.0 |
| *NDUFAF6* | 610 | 53.6 | 46.4 | 0 | 143 | 0 | 100 | 0 | 963 | 0 | 2.6 | 97.4 |
| *OTC* | 412 | 100 | 0 | 0 | 781 | 7.2 | 92.3 | 0.5 | 237 | 0 | 92.8 | 7.2 |
| *PALB2* | 976 | 90.7 | 9.3 | 0 | 846 | 0.1 | 39.2 | 60.6 | 6,692 | 0 | 0 | 100 |
| *PTEN* | 894 | 79.3 | 20.7 | 0 | 136 | 2.2 | 77.2 | 20.6 | 2,275 | 0 | 19.5 | 80.5 |
| *RAD51C* | 2,378 | 93.3 | 6.7 | 0 | 42 | 0 | 0 | 100 | 4,461 | 0 | 0.4 | 99.6 |
| *RAD51D* | 501 | 83.2 | 16.8 | 0 | 137 | 0.7 | 99.3 | 0 | 3,720 | 0 | 6.9 | 93.1 |
| *RHO* | 95 | 86.3 | 13.7 | 0 | 18 | 5.6 | 94.4 | 0 | 69 | 0 | 100 | 0 |
| *SCN5A* | 49 | 34.7 | 65.3 | 0 | 9 | 0 | 100 | 0 | 67 | 0 | 65.7 | 34.3 |
| *TSC2* | 771 | 38.4 | 61.6 | 0 | 186 | 0 | 100 | 0 | 2,063 | 0 | 99.6 | 0.4 |
| *VHL* | 327 | 76.8 | 23.2 | 0 | 241 | 2.5 | 76.8 | 20.7 | 1,700 | 0 | 5.3 | 94.7 |
| **All** | 29,264 | 75.8 | 21.9 | 2.3 | 11,310 | 4.6 | 79.0 | 16.4 | 88,620 | 0.3 | 32.0 | 67.7 |

Table S1: Gene-wise distribution of out-of-bag evidence direction assigned by ExCALIBR for variants with author-provided functional annotations. For each gene and author category (Abnormal, Indeterminate, Normal), the percentage of variants receiving pathogenic ($> 0$), indeterminate ($= 0$), or benign ($< 0$) evidence points is shown, alongside the total number of variants ($n$).

| Variant Group | P/LP | | | | VUS | | | | gnomAD | | | | B/LB | | | |
|---|---|---|---|---|---|---|---|---|---|---|---|---|---|---|---|---|
| Gene | $n$ | Path. | Ind. | Ben. | $n$ | Path. | Ind. | Ben. | $n$ | Path. | Ind. | Ben. | $n$ | Path. | Ind. | Ben. |
| *ASPA* | 128 | 28.1 | 71.9 | 0 | 114 | 7.9 | 92.1 | 0 | 704 | 13.2 | 86.8 | 0 | 12 | 0 | 100 | 0 |
| *BAP1* | 197 | 99.0 | 0.5 | 0.5 | 1,179 | 11.5 | 1.4 | 87.1 | 1,755 | 3.9 | 0.8 | 95.3 | 1,057 | 2.3 | 1.0 | 96.7 |
| *BARD1* | 184 | 98.9 | 0.5 | 0.5 | 1,661 | 14.9 | 2.6 | 82.4 | 1,927 | 9.9 | 1.6 | 88.5 | 959 | 0.2 | 1.0 | 98.7 |
| *BRCA1* | 319 | 93.7 | 4.1 | 2.2 | 656 | 19.2 | 9.9 | 70.9 | 970 | 18.2 | 5.6 | 76.2 | 230 | 2.2 | 2.6 | 95.2 |
| *BRCA2* | 513 | 77.4 | 17.0 | 5.7 | 1,764 | 14.7 | 11.8 | 73.4 | 1,820 | 15.1 | 10.1 | 74.8 | 854 | 4.4 | 9.0 | 86.5 |
| *CALM1* | 12 | 25.0 | 75.0 | 0 | 21 | 14.3 | 85.7 | 0 | 73 | 5.5 | 94.5 | 0 | 34 | 5.9 | 94.1 | 0 |
| *CARD11* | 8 | 0 | 100 | 0 | 60 | 0 | 100 | 0 | 148 | 0 | 100 | 0 | 14 | 0 | 100 | 0 |
| *CBS* | 38 | 21.1 | 78.9 | 0 | 172 | 2.3 | 97.7 | 0 | 75 | 4.0 | 96.0 | 0 | 282 | 0.7 | 99.3 | 0 |
| *CHEK2* | 103 | 66.0 | 24.3 | 9.7 | 1,344 | 25.5 | 19.6 | 54.9 | 792 | 22.0 | 20.6 | 57.4 | 236 | 4.2 | 16.9 | 78.8 |
| *CRX* | 12 | 0 | 100 | 0 | 113 | 0 | 100 | 0 | 349 | 0 | 100 | 0 | 8 | 0 | 100 | 0 |
| *CTCF* | 39 | 38.5 | 61.5 | 0 | 54 | 13.0 | 87.0 | 0 | 720 | 3.6 | 96.4 | 0 | 26 | 0 | 100 | 0 |
| *DDX3X* | 202 | 80.7 | 19.3 | 0 | 157 | 10.2 | 89.8 | 0 | 727 | 0.3 | 99.7 | 0 | 169 | 0 | 100 | 0 |
| *F9* | 575 | 50.4 | 29.9 | 19.7 | 275 | 26.9 | 48.4 | 24.7 | 1,825 | 5.4 | 40.3 | 54.2 | 700 | 1.0 | 33.0 | 66.0 |
| *FKRP* | 90 | 90.0 | 5.6 | 4.4 | 389 | 22.4 | 18.0 | 59.6 | 1,242 | 20.3 | 15.5 | 64.2 | 368 | 1.9 | 9.5 | 88.6 |
| *G6PD* | 189 | 24.3 | 75.7 | 0 | 103 | 7.8 | 92.2 | 0 | 457 | 2.8 | 97.2 | 0 | 106 | 0 | 100 | 0 |
| *GCK* | 596 | 57.4 | 42.6 | 0 | 370 | 36.2 | 63.8 | 0 | 1,188 | 17.3 | 82.7 | 0 | 85 | 1.2 | 98.8 | 0 |
| *HMBS* | 128 | 75.8 | 14.8 | 9.4 | 412 | 14.8 | 26.9 | 58.3 | 1,594 | 13.7 | 21.7 | 64.6 | 184 | 3.8 | 11.4 | 84.8 |
| *JAG1* | 16 | 0 | 100 | 0 | 160 | 15.0 | 77.5 | 7.5 | 459 | 0 | 91.3 | 8.7 | 99 | 0 | 93.9 | 6.1 |
| *KCNE1* | 21 | 66.7 | 33.3 | 0 | 287 | 12.2 | 43.2 | 44.6 | 204 | 14.2 | 46.1 | 39.7 | 81 | 1.2 | 59.3 | 39.5 |
| *KCNH2* | 222 | 86.0 | 8.6 | 5.4 | 999 | 17.2 | 8.1 | 74.7 | 1,751 | 9.1 | 8.2 | 82.6 | 375 | 6.1 | 5.1 | 88.8 |
| *KCNQ4* | 15 | 86.7 | 6.7 | 6.7 | 247 | 13.0 | 44.1 | 42.9 | 1,635 | 8.0 | 47.6 | 44.3 | 38 | 0 | 50.0 | 50.0 |
| *LARGE1* | 9 | 100 | 0 | 0 | 248 | 13.3 | 10.1 | 76.6 | 1,356 | 11.5 | 12.5 | 76.0 | 273 | 1.1 | 8.1 | 90.8 |
| *MSH2* | 44 | 86.4 | 6.8 | 6.8 | 1,425 | 3.3 | 4.4 | 92.4 | 1,353 | 1.7 | 3.8 | 94.5 | 19 | 5.3 | 0 | 94.7 |
| *NDUFAF6* | 6 | 50.0 | 50.0 | 0 | 55 | 12.7 | 30.9 | 56.4 | 347 | 15.3 | 26.5 | 58.2 | 6 | 0 | 16.7 | 83.3 |
| *OTC* | 79 | 65.8 | 32.9 | 1.3 | 80 | 31.2 | 63.7 | 5.0 | 139 | 13.7 | 80.6 | 5.8 | 7 | 0 | 71.4 | 28.6 |
| *PALB2* | 273 | 95.6 | 4.0 | 0.4 | 1,730 | 9.8 | 5.7 | 84.5 | 1,663 | 9.0 | 5.7 | 85.3 | 755 | 0.8 | 5.3 | 93.9 |
| *PAX6* | 192 | 91.7 | 6.8 | 1.6 | 174 | 55.2 | 10.9 | 33.9 | 384 | 10.2 | 18.8 | 71.1 | 4 | 0 | 0 | 100 |
| *PTEN* | 171 | 78.9 | 8.2 | 12.9 | 781 | 17.0 | 23.7 | 59.3 | 371 | 13.2 | 17.0 | 69.8 | 43 | 2.3 | 2.3 | 95.3 |
| *RAD51C* | 199 | 97.5 | 1.5 | 1.0 | 868 | 27.4 | 3.9 | 68.7 | 951 | 20.5 | 2.6 | 76.9 | 466 | 1.5 | 1.5 | 97.0 |
| *RAD51D* | 73 | 87.7 | 11.0 | 1.4 | 704 | 10.2 | 13.8 | 76.0 | 823 | 8.6 | 8.5 | 82.9 | 434 | 1.2 | 6.9 | 91.9 |
| *RHO* | 84 | 66.7 | 33.3 | 0 | 17 | 17.6 | 82.4 | 0 | 71 | 31.0 | 69.0 | 0 | 6 | 0 | 100 | 0 |
| *SCN5A* | 23 | 56.5 | 43.5 | 0 | 6 | 16.7 | 83.3 | 0 | 59 | 20.3 | 40.7 | 39.0 | 16 | 0 | 31.2 | 68.8 |
| *SGCB* | 12 | 83.3 | 8.3 | 8.3 | 121 | 6.6 | 33.1 | 60.3 | 443 | 12.2 | 26.4 | 61.4 | 0 | 0 | 0 | 0 |
| *TARDBP* | 12 | 0 | 100 | 0 | 19 | 0 | 89.5 | 10.5 | 61 | 0 | 90.2 | 9.8 | 0 | 0 | 0 | 0 |
| *TP53* | 1,627 | 83.6 | 8.9 | 7.5 | 7,328 | 22.8 | 17.2 | 59.9 | 7,384 | 22.3 | 9.5 | 68.1 | 727 | 4.7 | 10.7 | 84.6 |
| *TPK1* | 8 | 37.5 | 62.5 | 0 | 93 | 29.0 | 66.7 | 4.3 | 395 | 26.3 | 73.7 | 0 | 45 | 6.7 | 93.3 | 0 |
| *TSC2* | 61 | 63.9 | 36.1 | 0 | 848 | 6.1 | 93.8 | 0.1 | 937 | 2.6 | 96.7 | 0.7 | 344 | 0.3 | 98.5 | 1.2 |
| *VHL* | 199 | 71.4 | 18.6 | 10.1 | 418 | 2.9 | 17.5 | 79.7 | 488 | 2.5 | 14.1 | 83.4 | 236 | 1.3 | 8.5 | 90.3 |
| *XRCC2* | 6 | 33.3 | 0 | 66.7 | 338 | 0.9 | 0 | 99.1 | 621 | 1.6 | 0.3 | 98.1 | 163 | 0 | 0 | 100 |
| **All** | 6,685 | 74.7 | 19.7 | 5.5 | 25,790 | 17.0 | 20.1 | 62.9 | 38,261 | 12.5 | 26.7 | 60.8 | 9,461 | 2.0 | 20.5 | 77.5 |

Table S2: Gene-wise distribution of out-of-bag evidence direction assigned by ExCALIBR for variants stratified by variant group. For each gene and variant group (P/LP, VUS, gnomAD, B/LB), the percentage of variants receiving pathogenic ($> 0$), indeterminate ($= 0$), or benign ($< 0$) evidence points is shown, alongside the total number of variants ($n$).

Table S3: Performance comparison of out-of-bag ExCALIBR-calibrated evidence vs. author-provided functional annotations on P/LP and B/LB variants from ClinVar. Pathogenic and benign refer to the direction of evidence assigned.

| Metric | Calibrated Evidence | Author Annotations |
|---|---|---|
| Total variants | 9,772 | 9,772 |
| Determinate assignments | 7,846 (80.3%) | 9,177 (93.9%) |
| Indeterminate assignments | 1,926 (19.7%) | 595 (6.1%) |
| Accuracy | 0.979 | 0.936 |
| Sensitivity | 0.972 | 0.882 |
| Specificity | 0.982 | 0.966 |
| MCC | 0.953 | 0.859 |
| $LR^+$ (pathogenic vs. benign) | 55.5 | 25.8 |
| $LR^+$ (pathogenic vs. rest) | 52.3 | 24.9 |
| $LR^+$ (benign vs. rest) | 37.1 | 8.5 |
| DOR (pathogenic vs. benign) | 1941.7 | 210.6 |
| DOR (pathogenic vs. rest) | 206.8 | 126.4 |
| DOR (benign vs. rest) | 186.8 | 92.4 |

Table S4: Performance comparison of out-of-bag ExCALIBR-calibrated evidence vs. author-provided functional annotations on P/LP and B/LB variants from the ClinGen Evidence Repository. Pathogenic and benign refer to the direction of evidence assigned. $LR^+$ and DOR values of $\infty$ are infinite where no benign variants were misclassified as pathogenic; misclassifications among pathogenic variants are still present.

| Metric | Calibrated Evidence | Author Annotations |
|---|---|---|
| Total variants | 276 | 276 |
| Determinate assignments | 160 (58.0%) | 215 (77.9%) |
| Indeterminate assignments | 116 (42.0%) | 61 (22.1%) |
| Accuracy | 0.938 | 0.874 |
| Sensitivity | 0.934 | 0.871 |
| Specificity | 1.000 | 0.929 |
| MCC | 0.665 | 0.512 |
| $LR^+$ (pathogenic vs. benign) | $\infty$ | 12.2 |
| $LR^+$ (pathogenic vs. rest) | $\infty$ | 14.4 |
| $LR^+$ (benign vs. rest) | 10.9 | 6.1 |
| DOR (pathogenic vs. benign) | $\infty$ | 87.5 |
| DOR (pathogenic vs. rest) | $\infty$ | 43.8 |
| DOR (benign vs. rest) | 18.4 | 14.3 |

| Alternate parameterization | | Related quantities |
|---|---|---|
| Canonical →Alternate | Alternate → Canonical | |
| $\Delta = \omega\delta$ | $\lambda = \mathrm{sign}(\Delta)\sqrt{\Delta^2/\Gamma}$ | $\delta = \lambda/\sqrt{1+\lambda^2}$ |
| $\Gamma = \omega^2 - \Delta^2$ | $\omega = \sqrt{\Gamma + \Delta^2}$ | |

Table S5: Parameterizations of the skew normal distribution

$$\bar{m}^{(*)}(s, \Delta) = s - \nu(s, \bar{\theta}^{(*)}) \cdot \Delta$$
$$\bar{d}^{(*)}(s, \mu) = \nu(s, \bar{\theta}^{(*)}) \cdot (s - \mu)$$
$$\bar{g}^{(*)}(s, \mu, \Delta) = (s - \mu)^2 - 2 \cdot \Delta \cdot \nu(s, \bar{\theta}^{(*)}) \cdot (s - \mu) + \Delta^2 \cdot w(s, \bar{\theta}^{(*)})$$
$$\nu(s, \theta) = \mathbb{E}[T_s]$$
$$w(s, \theta) = \mathbb{E}[T_s^2]$$
$$T_s \sim \mathrm{TN}\left(\frac{\delta}{\omega}(s - \mu), 1 - \delta^2, \mathbb{R}^+\right)$$

Table S6: Related quantities for EM updates

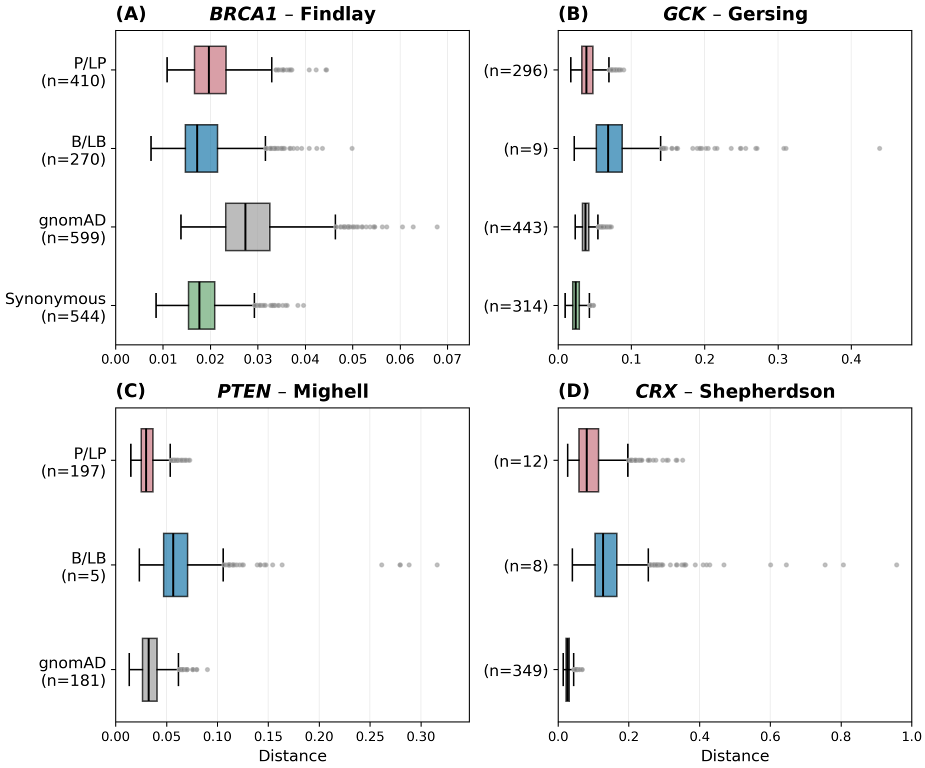

Figure S1: Distributions of normalized distances between empirical and estimated cumulative distribution functions for each sample, obtained using 1,000 bootstrapping iterations, for the same datasets as in Figure 2. Datasets for panels A, B, C, and D were taken from Findlay *et al.* [7], Gersing *et al.* [8], Mighell *et al.* [9], and Shepherdson *et al.* [10], respectively.

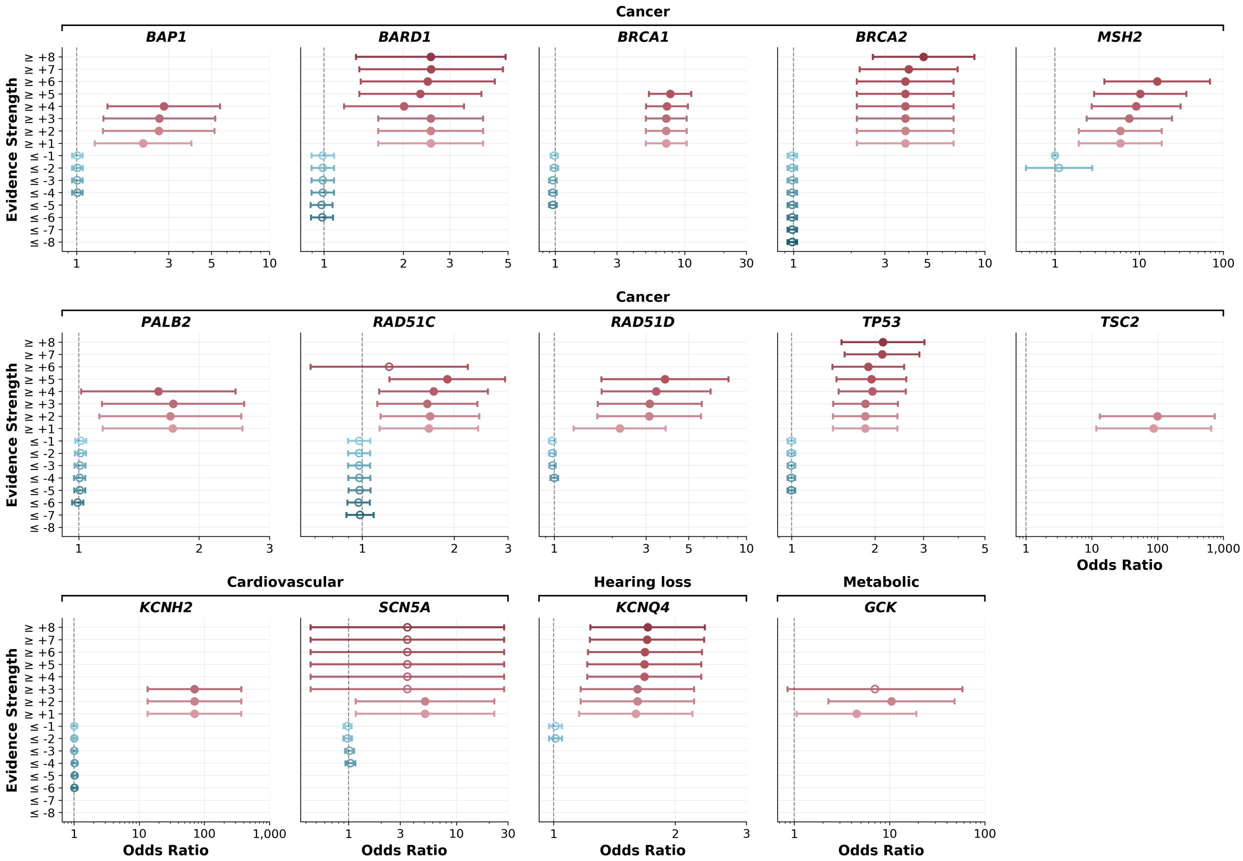

Figure S2: Odds ratios for occurrence of disease in individuals with variants meeting each ExCALIBR-calibrated evidence strength threshold in the All of Us biobank for genes that had significant association with disease at some level of pathogenic evidence. Genes are stratified by the type of disease association. Circles denote the estimated odds ratio and whiskers denote the 95% confidence interval. Filled circles denote estimates whose 95% confidence interval does not include 1; open circles denote those whose interval includes 1. The x-axes show odds ratios on a log scale.

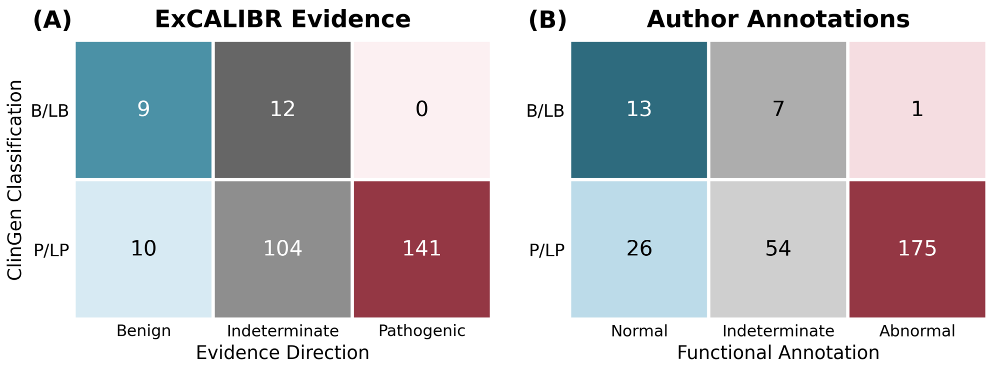

Figure S3: Comparison of ExCALIBR out-of-bag evidence assignments (A) and author-provided annotations (B) for ClinGen Evidence Repository P/LP and B/LB variants in author-annotated datasets. Evidence assignments less than zero indicate benign direction; greater than zero indicate pathogenic direction; zero is indeterminate.
