## Supplementary material for "Gene-based calibration of high-throughput functional assays for clinical variant classification": Model Fit Visualizations

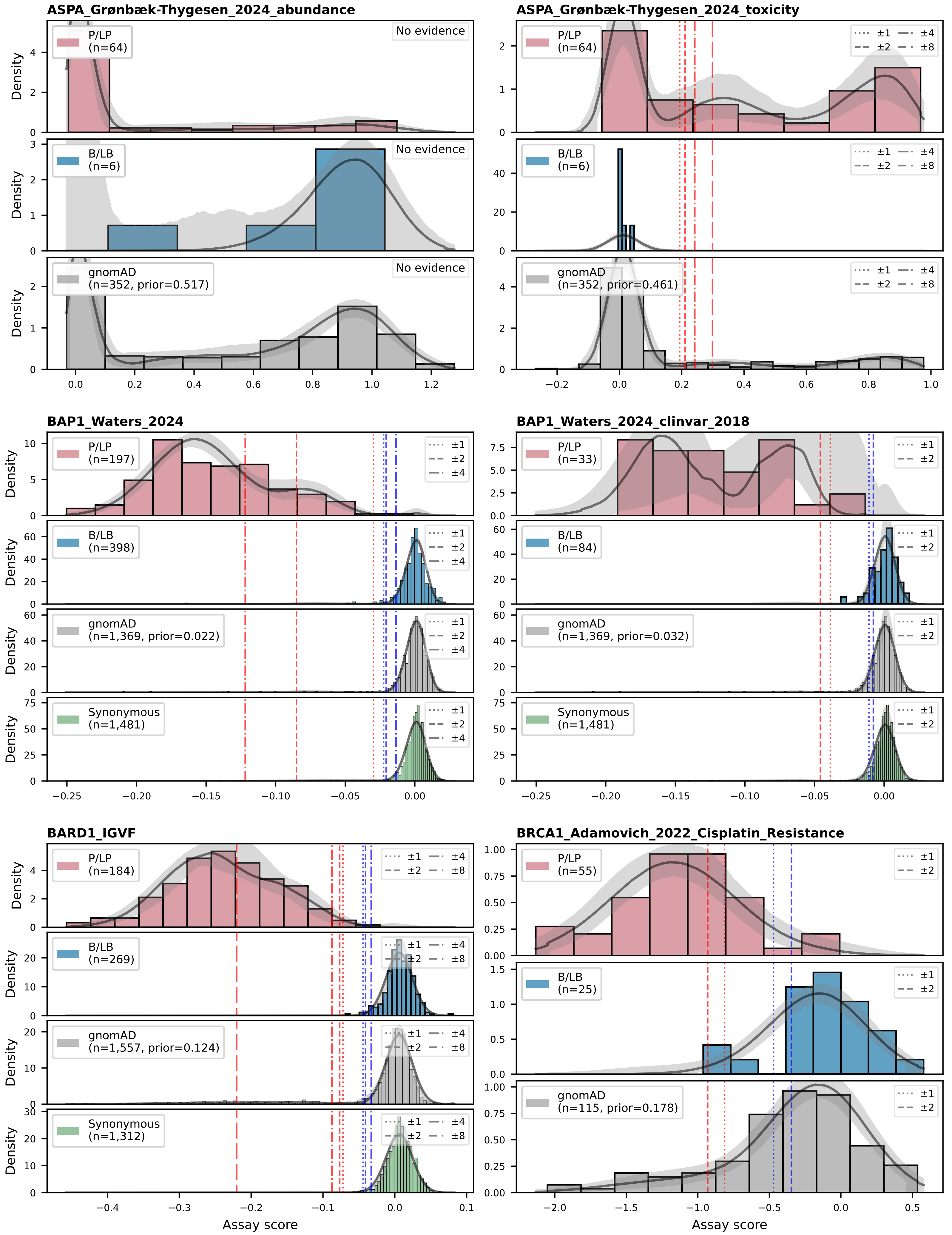
ASPA_Grønbæk-Thygesen_2024_abundance
P/LP (n=64)
B/LB (n=6)
gnomAD (n=352, prior=0.517)
No evidence
ASPA_Grønbæk-Thygesen_2024_toxicity
P/LP (n=64)
B/LB (n=6)
gnomAD (n=352, prior=0.461)
±1 ±2 ±4 ±8
BAP1_Waters_2024
P/LP (n=197)
B/LB (n=398)
gnomAD (n=1,369, prior=0.022)
Synonymous (n=1,481)
±1 ±2 ±4
BAP1_Waters_2024_clinvar_2018
P/LP (n=33)
B/LB (n=84)
gnomAD (n=1,369, prior=0.032)
Synonymous (n=1,481)
±1 ±2
BARD1_IGVF
P/LP (n=184)
B/LB (n=269)
gnomAD (n=1,557, prior=0.124)
Synonymous (n=1,312)
±1 ±2 ±4 ±8
BRCA1_Adamovich_2022_Cisplatin_Resistance
P/LP (n=55)
B/LB (n=25)
gnomAD (n=115, prior=0.178)
±1 ±2
Density
Assay score

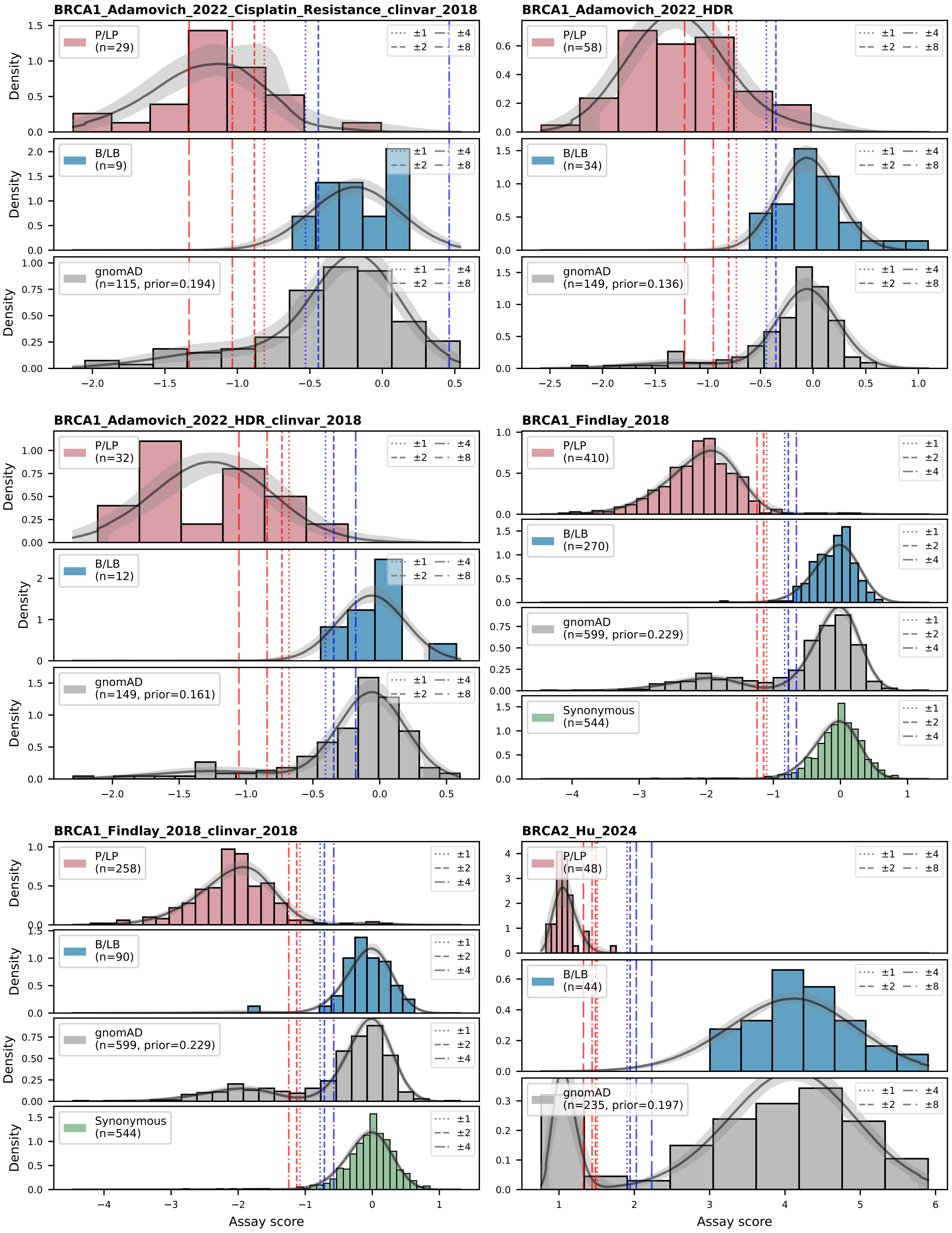

BRCA1_Adamovich_2022_Cisplatin_Resistance_clinvar_2018
P/LP (n=29)
B/LB (n=9)
gnomAD (n=115, prior=0.194)
BRCA1_Adamovich_2022_HDR
P/LP (n=58)
B/LB (n=34)
gnomAD (n=149, prior=0.136)
BRCA1_Adamovich_2022_HDR_clinvar_2018
P/LP (n=32)
B/LB (n=12)
gnomAD (n=149, prior=0.161)
BRCA1_Findlay_2018
P/LP (n=410)
B/LB (n=270)
gnomAD (n=599, prior=0.229)
Synonymous (n=544)
BRCA1_Findlay_2018_clinvar_2018
P/LP (n=258)
B/LB (n=90)
gnomAD (n=599, prior=0.229)
Synonymous (n=544)
BRCA2_Hu_2024
P/LP (n=48)
B/LB (n=44)
gnomAD (n=235, prior=0.197)
±1 ±2 ±4 ±8
Density
Assay score

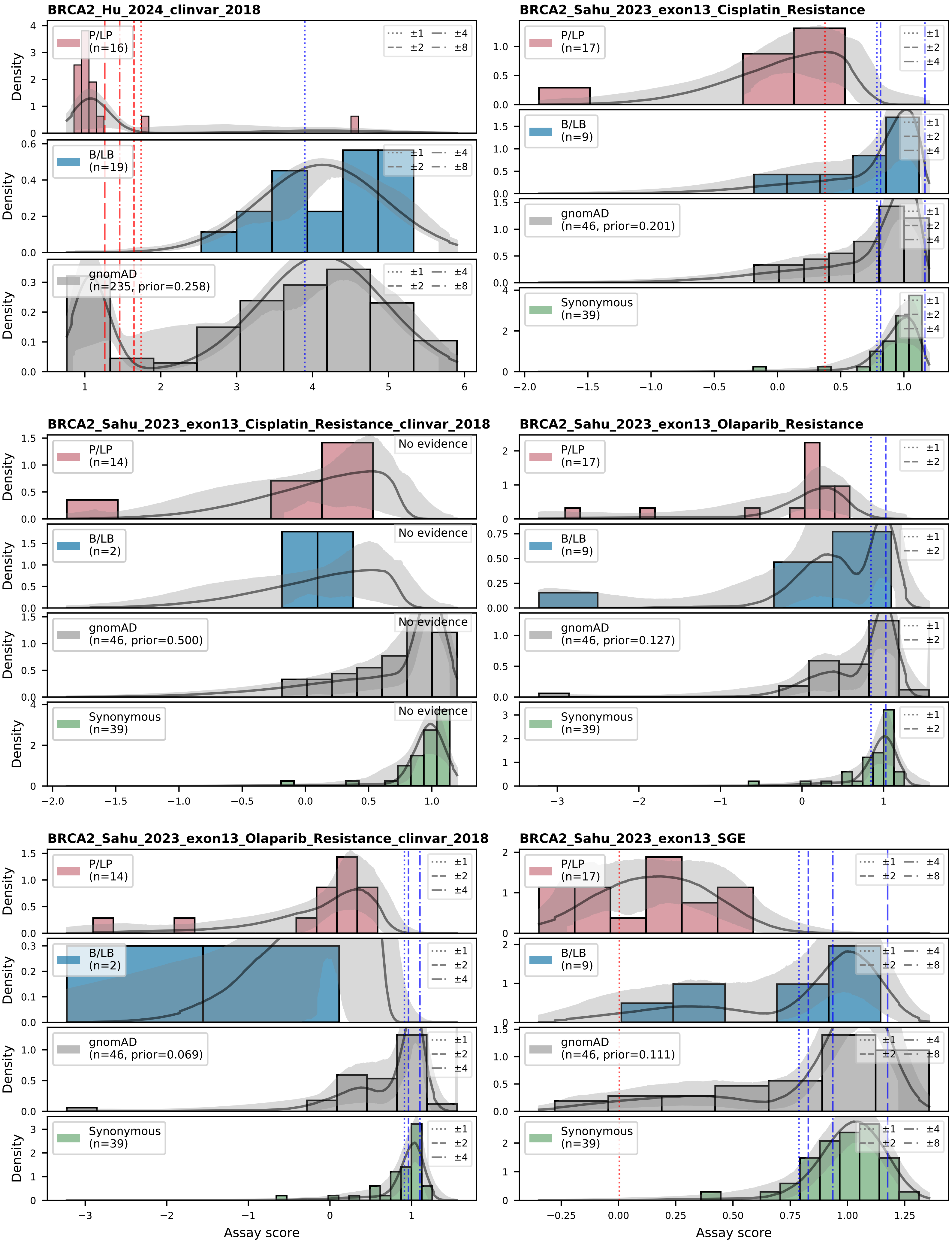
BRCA2_Hu_2024_clinvar_2018
P/LP (n=16)
B/LB (n=19)
gnomAD (n=235, prior=0.258)
±1 ±2 ±4 ±8
BRCA2_Sahu_2023_exon13_Cisplatin_Resistance
P/LP (n=17)
B/LB (n=9)
gnomAD (n=46, prior=0.201)
Synonymous (n=39)
±1 ±2 ±4
BRCA2_Sahu_2023_exon13_Cisplatin_Resistance_clinvar_2018
P/LP (n=14)
B/LB (n=2)
gnomAD (n=46, prior=0.500)
Synonymous (n=39)
No evidence
BRCA2_Sahu_2023_exon13_Olaparib_Resistance
P/LP (n=17)
B/LB (n=9)
gnomAD (n=46, prior=0.127)
Synonymous (n=39)
±1 ±2
BRCA2_Sahu_2023_exon13_Olaparib_Resistance_clinvar_2018
P/LP (n=14)
B/LB (n=2)
gnomAD (n=46, prior=0.069)
Synonymous (n=39)
±1 ±2 ±4
BRCA2_Sahu_2023_exon13_SGE
P/LP (n=17)
B/LB (n=9)
gnomAD (n=46, prior=0.111)
Synonymous (n=39)
±1 ±2 ±4 ±8
Density
Assay score

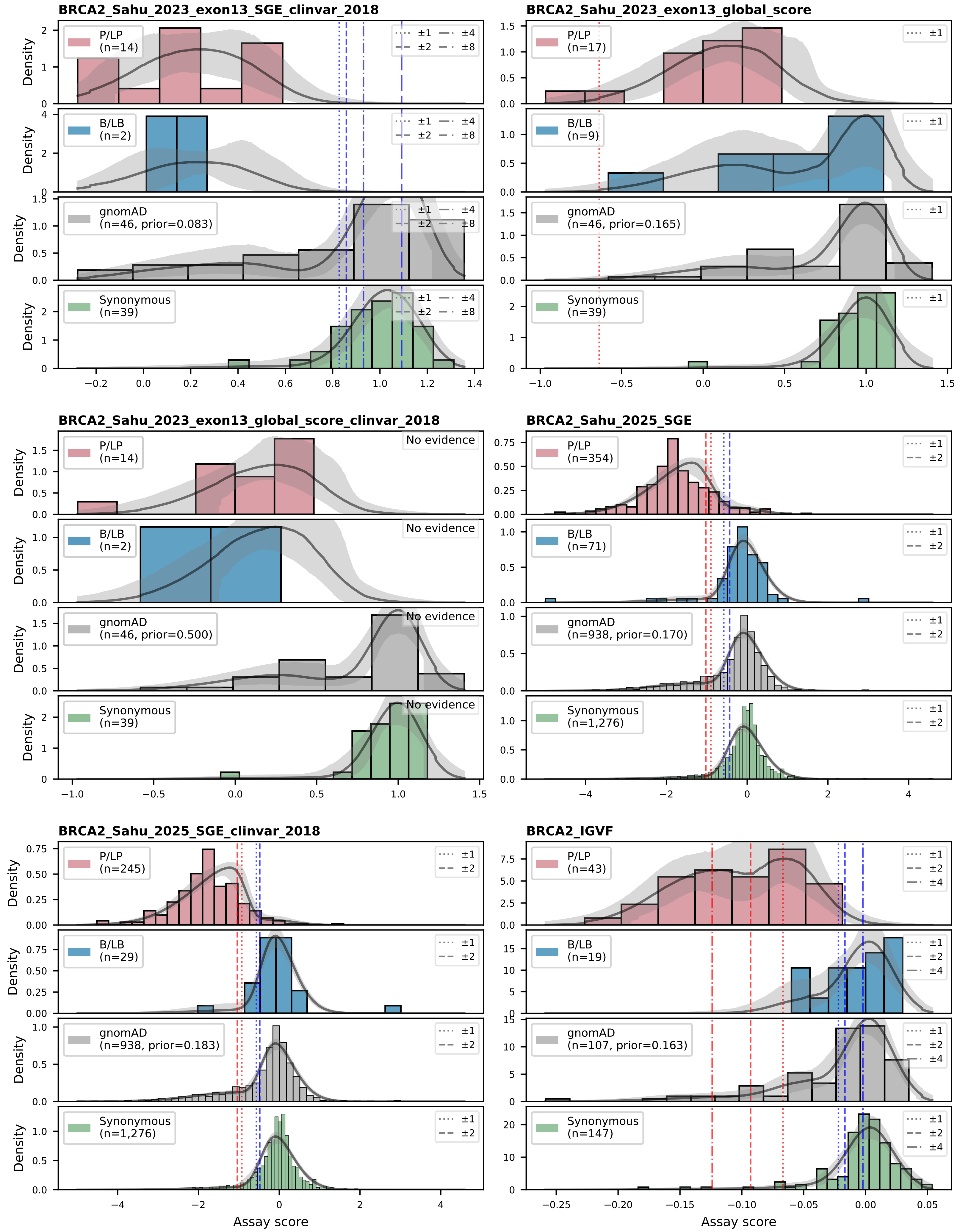
BRCA2_Sahu_2023_exon13_SGE_clinvar_2018
P/LP (n=14)
B/LB (n=2)
gnomAD (n=46, prior=0.083)
Synonymous (n=39)
±1 ±2 ±4 ±8
BRCA2_Sahu_2023_exon13_global_score
P/LP (n=17)
B/LB (n=9)
gnomAD (n=46, prior=0.165)
Synonymous (n=39)
±1
BRCA2_Sahu_2023_exon13_global_score_clinvar_2018
P/LP (n=14)
B/LB (n=2)
gnomAD (n=46, prior=0.500)
Synonymous (n=39)
No evidence
BRCA2_Sahu_2025_SGE
P/LP (n=354)
B/LB (n=71)
gnomAD (n=938, prior=0.170)
Synonymous (n=1,276)
±1 ±2
BRCA2_Sahu_2025_SGE_clinvar_2018
P/LP (n=245)
B/LB (n=29)
gnomAD (n=938, prior=0.183)
Synonymous (n=1,276)
±1 ±2
BRCA2_IGVF
P/LP (n=43)
B/LB (n=19)
gnomAD (n=107, prior=0.163)
Synonymous (n=147)
±1 ±2 ±4
Density
Assay score

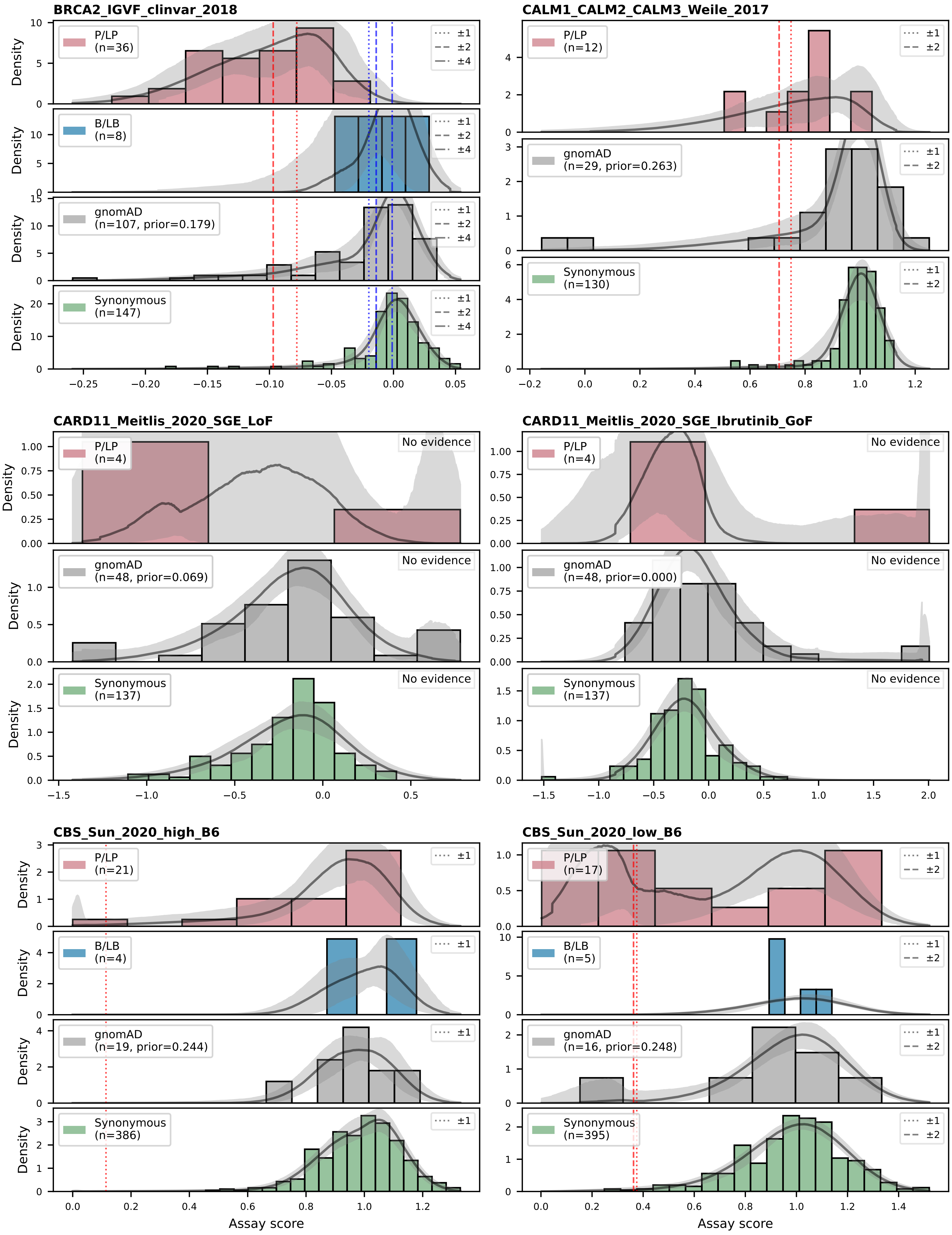
BRCA2_IGVF_clinvar_2018
P/LP (n=36)
B/LB (n=8)
gnomAD (n=107, prior=0.179)
Synonymous (n=147)
±1
±2
±4
CALM1_CALM2_CALM3_Weile_2017
P/LP (n=12)
gnomAD (n=29, prior=0.263)
Synonymous (n=130)
CARD11_Meitlis_2020_SGE_LoF
P/LP (n=4)
gnomAD (n=48, prior=0.069)
Synonymous (n=137)
No evidence
CARD11_Meitlis_2020_SGE_Ibrutinib_GoF
P/LP (n=4)
gnomAD (n=48, prior=0.000)
Synonymous (n=137)
CBS_Sun_2020_high_B6
P/LP (n=21)
B/LB (n=4)
gnomAD (n=19, prior=0.244)
Synonymous (n=386)
CBS_Sun_2020_low_B6
P/LP (n=17)
B/LB (n=5)
gnomAD (n=16, prior=0.248)
Synonymous (n=395)
Density
Assay score

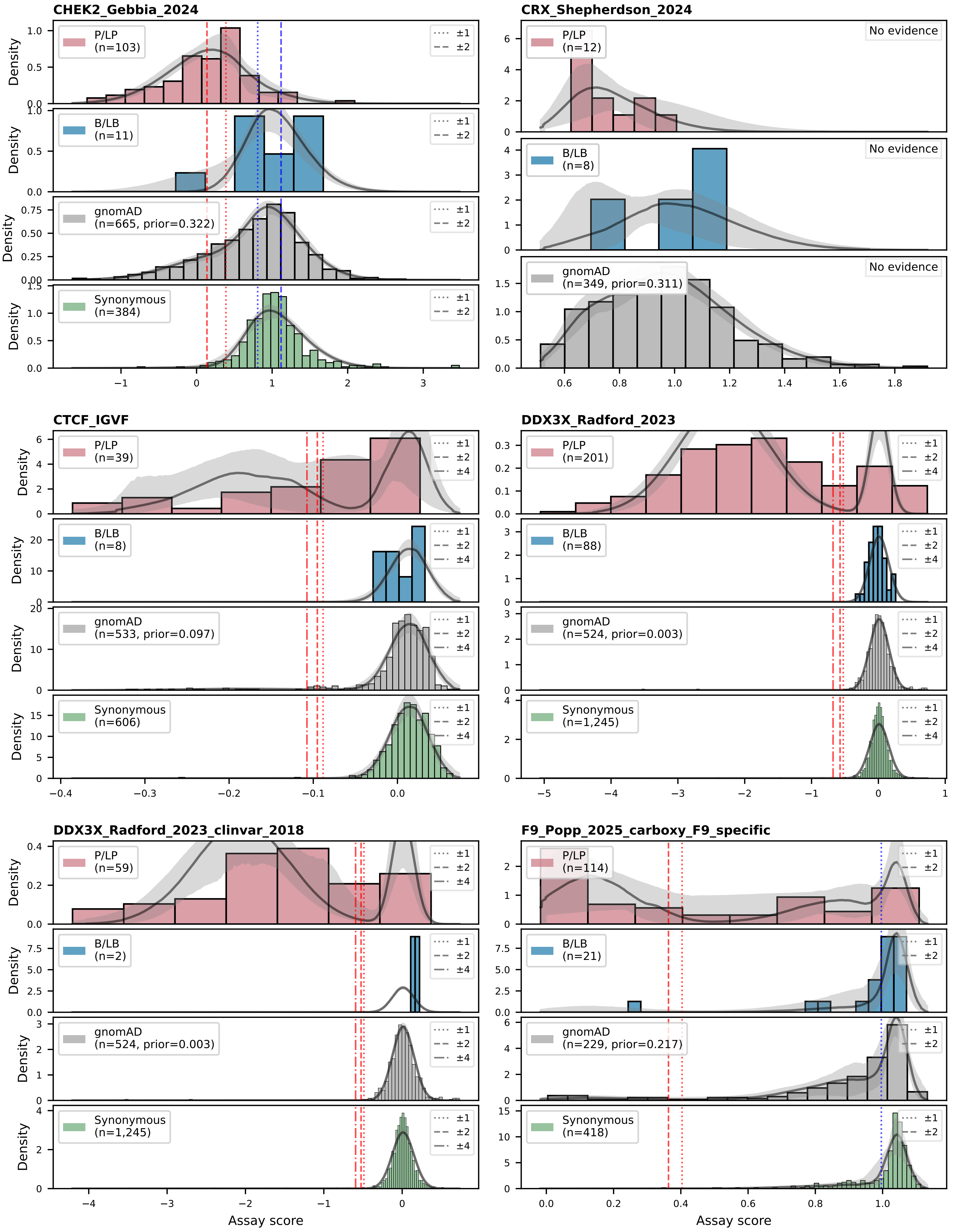

CHEK2_Gebbia_2024
P/LP (n=103)
B/LB (n=11)
gnomAD (n=665, prior=0.322)
Synonymous (n=384)
±1
±2
Density
CRX_Shepherdson_2024
No evidence
P/LP (n=12)
B/LB (n=8)
gnomAD (n=349, prior=0.311)
CTCF_IGVF
P/LP (n=39)
B/LB (n=8)
gnomAD (n=533, prior=0.097)
Synonymous (n=606)
±4
DDX3X_Radford_2023
P/LP (n=201)
B/LB (n=88)
gnomAD (n=524, prior=0.003)
Synonymous (n=1,245)
DDX3X_Radford_2023_clinvar_2018
P/LP (n=59)
B/LB (n=2)
gnomAD (n=524, prior=0.003)
Synonymous (n=1,245)
F9_Popp_2025_carboxy_F9_specific
P/LP (n=114)
B/LB (n=21)
gnomAD (n=229, prior=0.217)
Synonymous (n=418)
Assay score

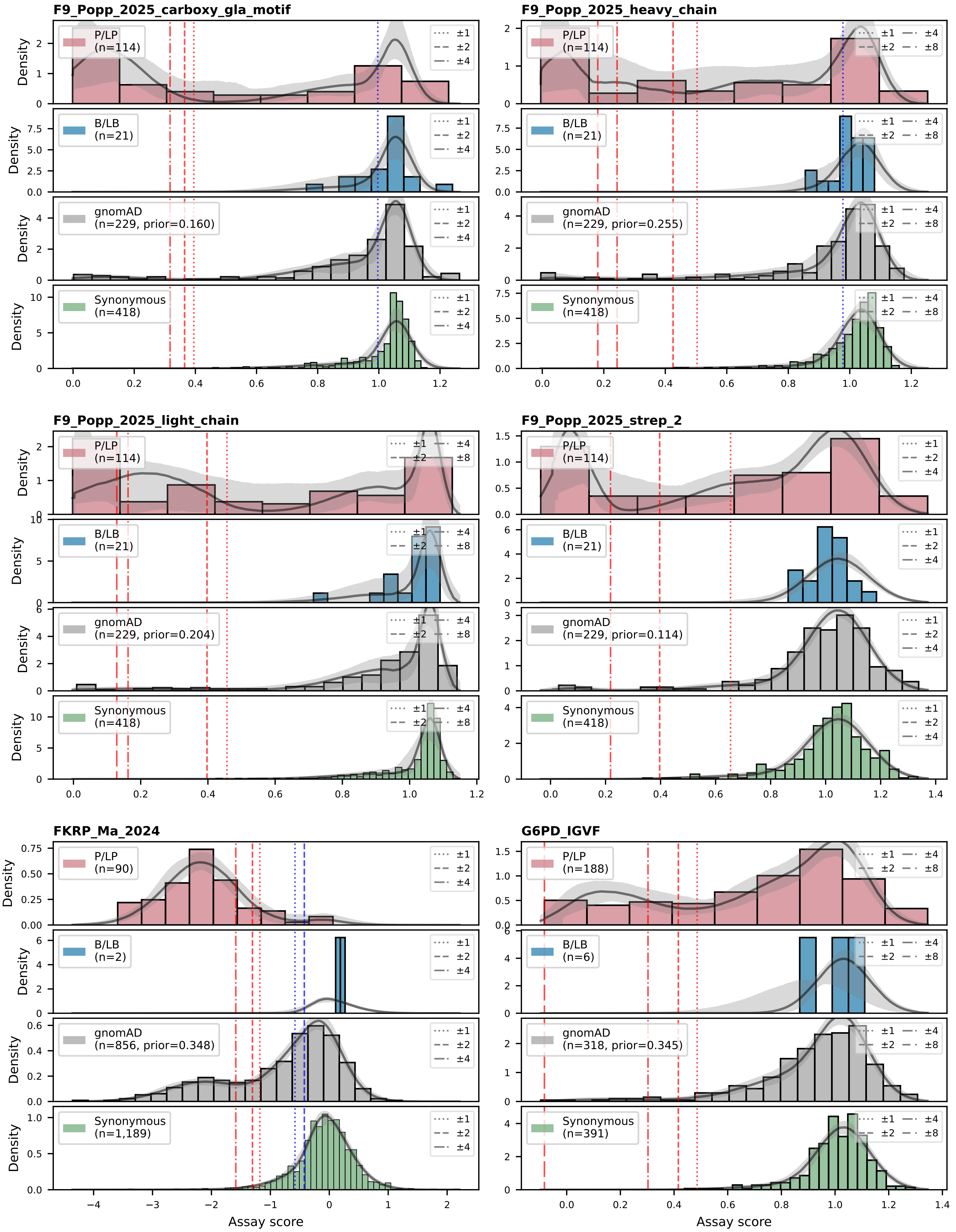
F9_Popp_2025_carboxy_gla_motif
F9_Popp_2025_heavy_chain
F9_Popp_2025_light_chain
F9_Popp_2025_strep_2
FKRP_Ma_2024
G6PD_IGVF
P/LP (n=114)
B/LB (n=21)
gnomAD (n=229, prior=0.160)
gnomAD (n=229, prior=0.255)
gnomAD (n=229, prior=0.204)
gnomAD (n=229, prior=0.114)
Synonymous (n=418)
P/LP (n=90)
B/LB (n=2)
gnomAD (n=856, prior=0.348)
Synonymous (n=1,189)
P/LP (n=188)
B/LB (n=6)
gnomAD (n=318, prior=0.345)
Synonymous (n=391)
±1
±2
±4
±8
Density
Assay score

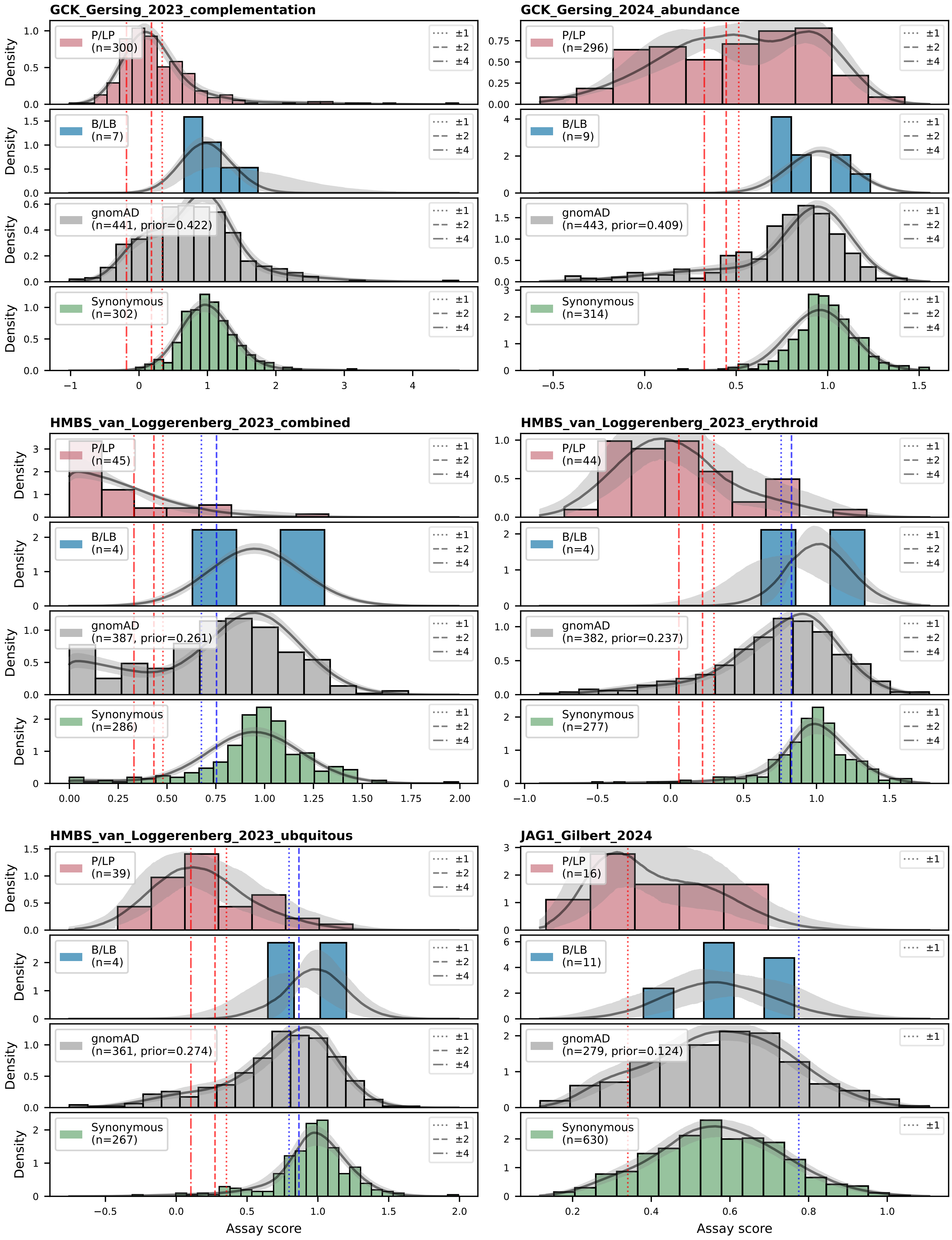
GCK_Gersing_2023_complementation
GCK_Gersing_2024_abundance
HMBS_van_Loggerenberg_2023_combined
HMBS_van_Loggerenberg_2023_erythroid
HMBS_van_Loggerenberg_2023_ubquitous
JAG1_Gilbert_2024
P/LP
(n=300)
B/LB
(n=7)
gnomAD
(n=441, prior=0.422)
Synonymous
(n=302)
P/LP
(n=296)
B/LB
(n=9)
gnomAD
(n=443, prior=0.409)
Synonymous
(n=314)
P/LP
(n=45)
B/LB
(n=4)
gnomAD
(n=387, prior=0.261)
Synonymous
(n=286)
P/LP
(n=44)
B/LB
(n=4)
gnomAD
(n=382, prior=0.237)
Synonymous
(n=277)
P/LP
(n=39)
B/LB
(n=4)
gnomAD
(n=361, prior=0.274)
Synonymous
(n=267)
P/LP
(n=16)
B/LB
(n=11)
gnomAD
(n=279, prior=0.124)
Synonymous
(n=630)
±1
±2
±4
Density
Assay score

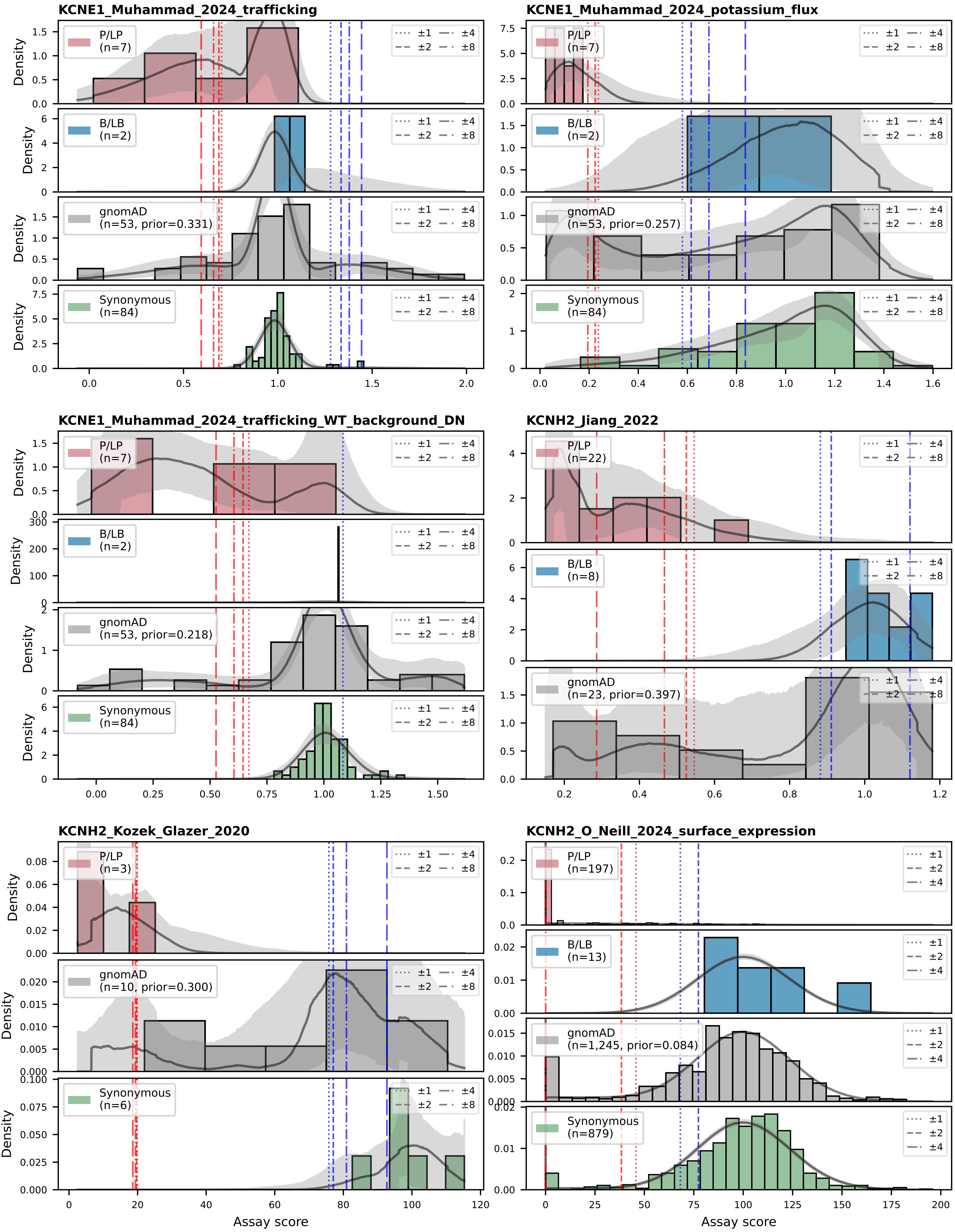
KCNE1_Muhammad_2024_trafficking
KCNE1_Muhammad_2024_potassium_flux
KCNE1_Muhammad_2024_trafficking_WT_background_DN
KCNH2_Jiang_2022
KCNH2_Kozek_Glazer_2020
KCNH2_O_Neill_2024_surface_expression
P/LP (n=7)
B/LB (n=2)
gnomAD (n=53, prior=0.331)
Synonymous (n=84)
gnomAD (n=53, prior=0.257)
gnomAD (n=53, prior=0.218)
P/LP (n=22)
B/LB (n=8)
gnomAD (n=23, prior=0.397)
P/LP (n=3)
gnomAD (n=10, prior=0.300)
Synonymous (n=6)
P/LP (n=197)
B/LB (n=13)
gnomAD (n=1,245, prior=0.084)
Synonymous (n=879)
±1
±2
±4
±8
Density
Assay score

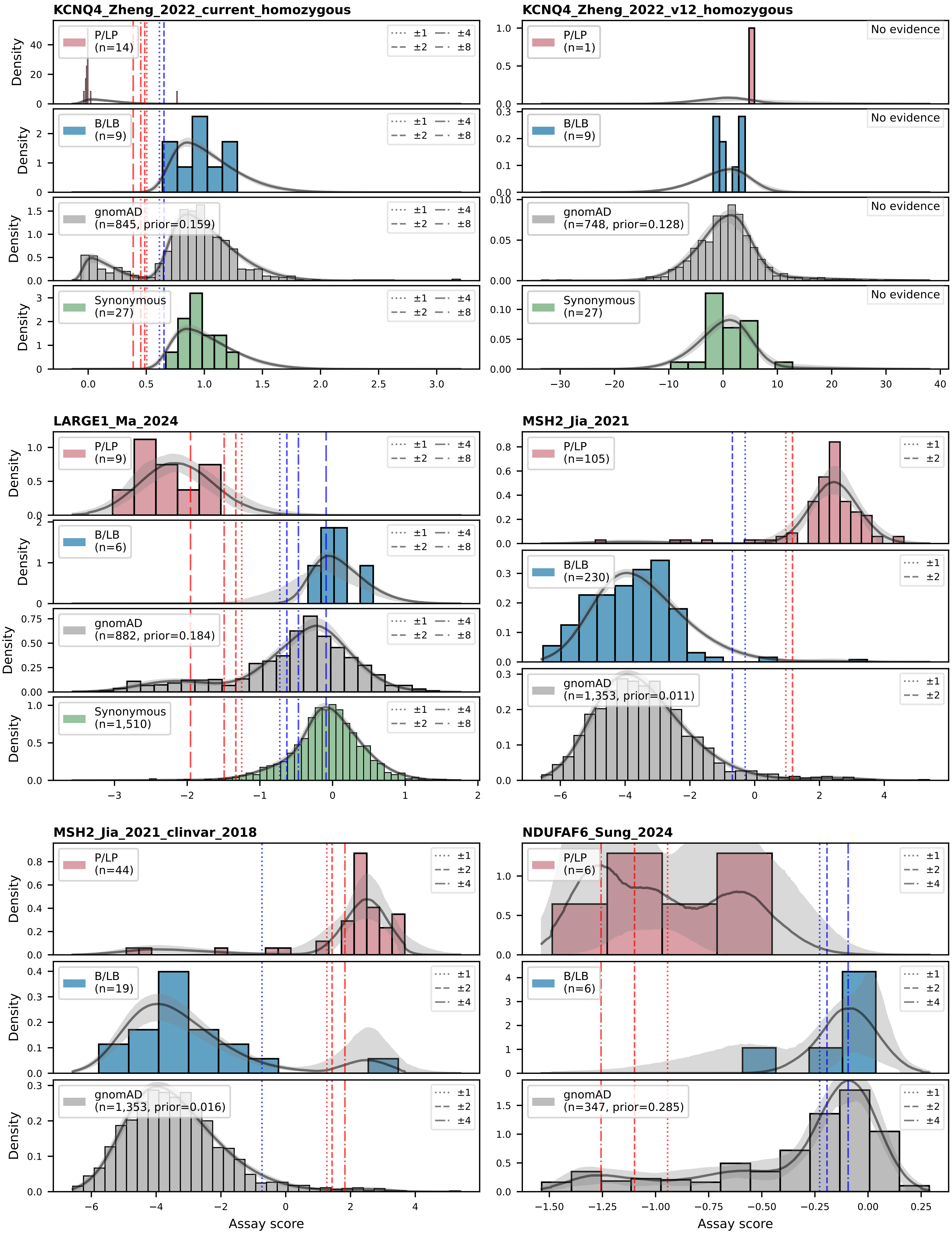
KCNQ4_Zheng_2022_current_homozygous
KCNQ4_Zheng_2022_v12_homozygous
LARGE1_Ma_2024
MSH2_Jia_2021
MSH2_Jia_2021_clinvar_2018
NDUFAF6_Sung_2024
P/LP
(n=14)
(n=1)
(n=9)
(n=105)
(n=44)
(n=6)
B/LB
(n=9)
(n=6)
(n=230)
(n=19)
gnomAD
(n=845, prior=0.159)
(n=748, prior=0.128)
(n=882, prior=0.184)
(n=1,353, prior=0.011)
(n=1,353, prior=0.016)
(n=347, prior=0.285)
Synonymous
(n=27)
(n=1,510)
±1
±2
±4
±8
No evidence
Density
Assay score

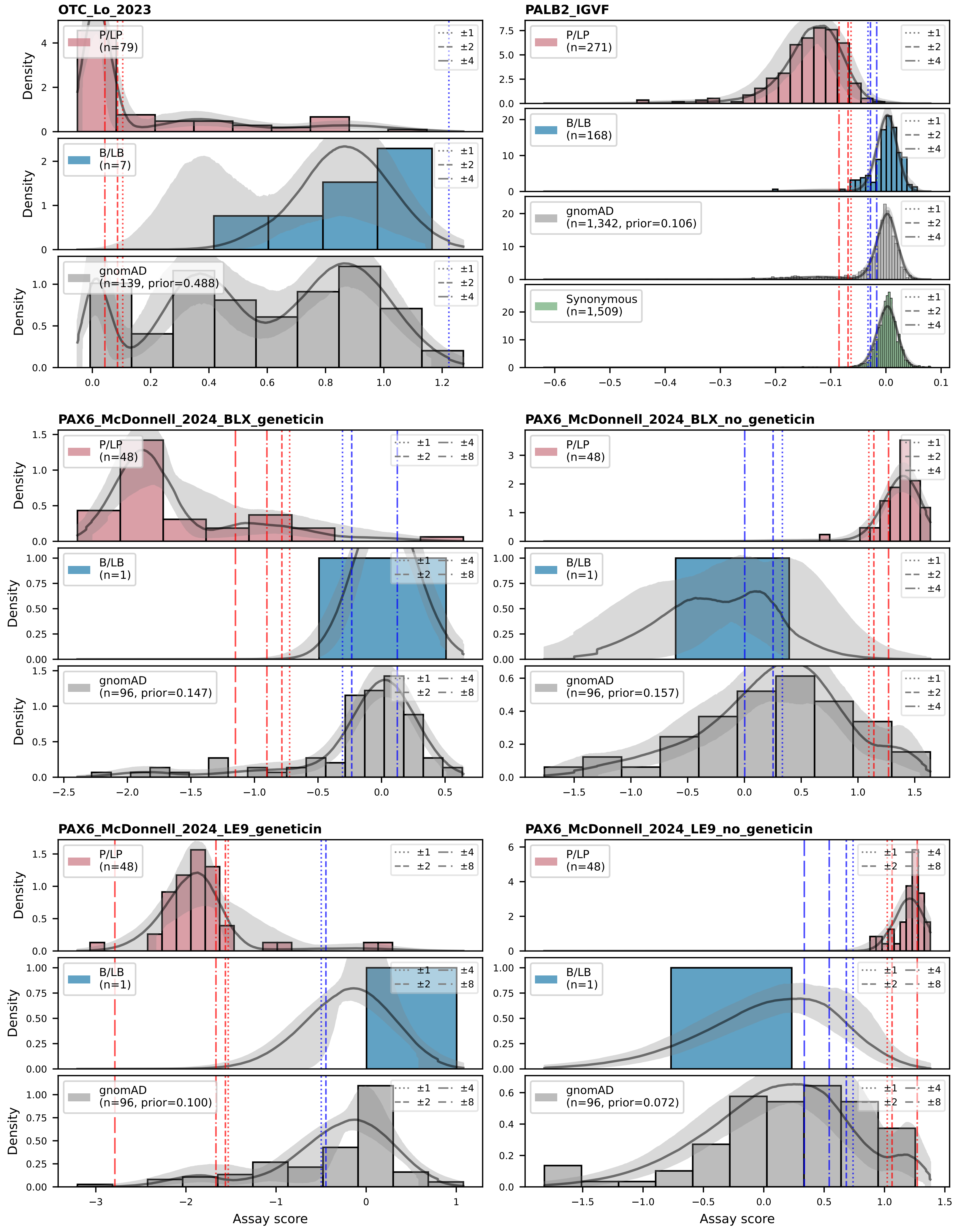
OTC_Lo_2023
PALB2_IGVF
P/LP (n=79)
B/LB (n=7)
gnomAD (n=139, prior=0.488)
P/LP (n=271)
B/LB (n=168)
gnomAD (n=1,342, prior=0.106)
Synonymous (n=1,509)
PAX6_McDonnell_2024_BLX_geneticin
PAX6_McDonnell_2024_BLX_no_geneticin
P/LP (n=48)
B/LB (n=1)
gnomAD (n=96, prior=0.147)
gnomAD (n=96, prior=0.157)
PAX6_McDonnell_2024_LE9_geneticin
PAX6_McDonnell_2024_LE9_no_geneticin
gnomAD (n=96, prior=0.100)
gnomAD (n=96, prior=0.072)
±1
±2
±4
±8
Density
Assay score

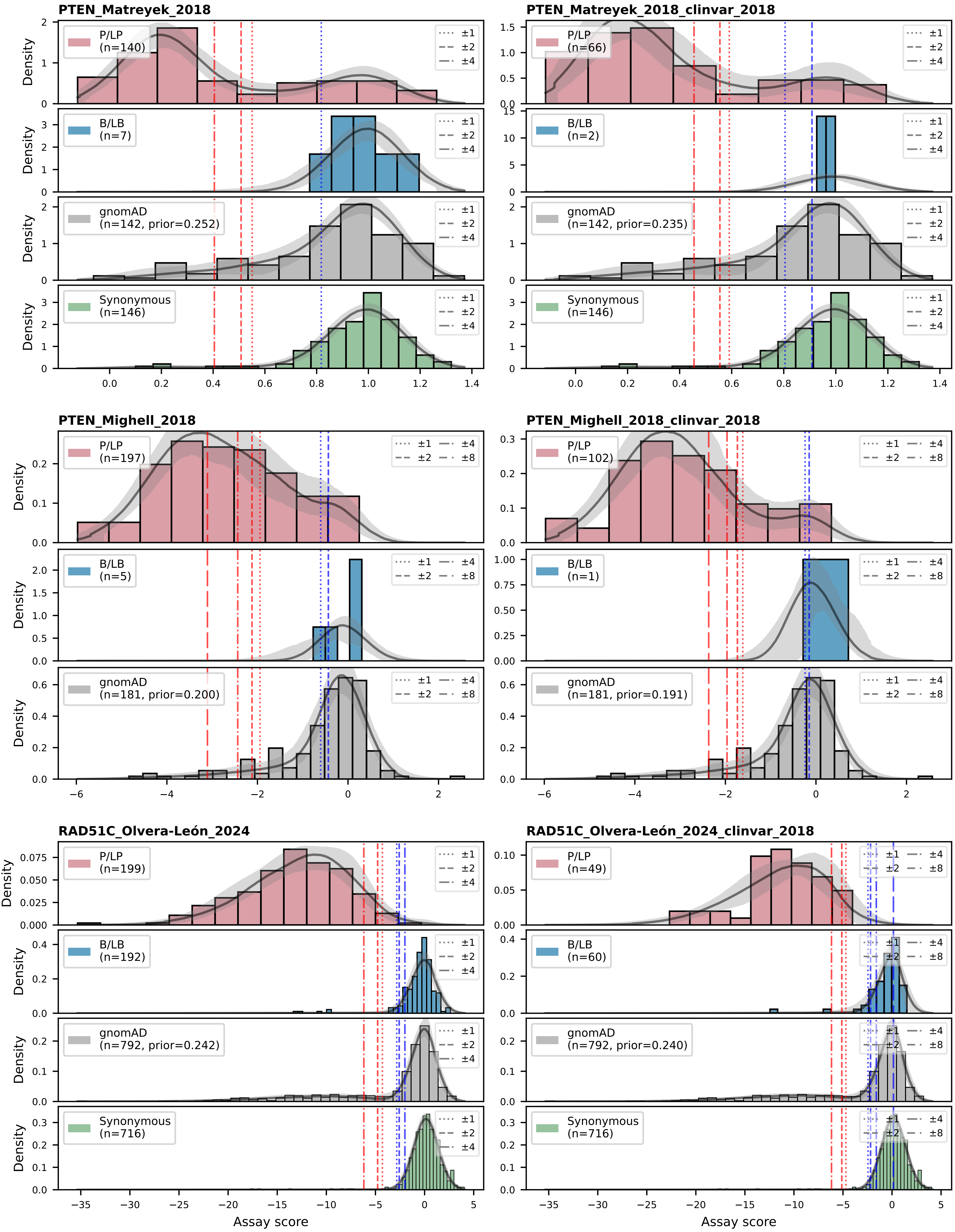
PTEN_Matreyek_2018
P/LP (n=140)
B/LB (n=7)
gnomAD (n=142, prior=0.252)
Synonymous (n=146)
PTEN_Matreyek_2018_clinvar_2018
P/LP (n=66)
B/LB (n=2)
gnomAD (n=142, prior=0.235)
Synonymous (n=146)
±1
±2
±4
PTEN_Mighell_2018
P/LP (n=197)
B/LB (n=5)
gnomAD (n=181, prior=0.200)
PTEN_Mighell_2018_clinvar_2018
P/LP (n=102)
B/LB (n=1)
gnomAD (n=181, prior=0.191)
±8
RAD51C_Olvera-León_2024
P/LP (n=199)
B/LB (n=192)
gnomAD (n=792, prior=0.242)
Synonymous (n=716)
RAD51C_Olvera-León_2024_clinvar_2018
P/LP (n=49)
B/LB (n=60)
gnomAD (n=792, prior=0.240)
Synonymous (n=716)
Density
Assay score

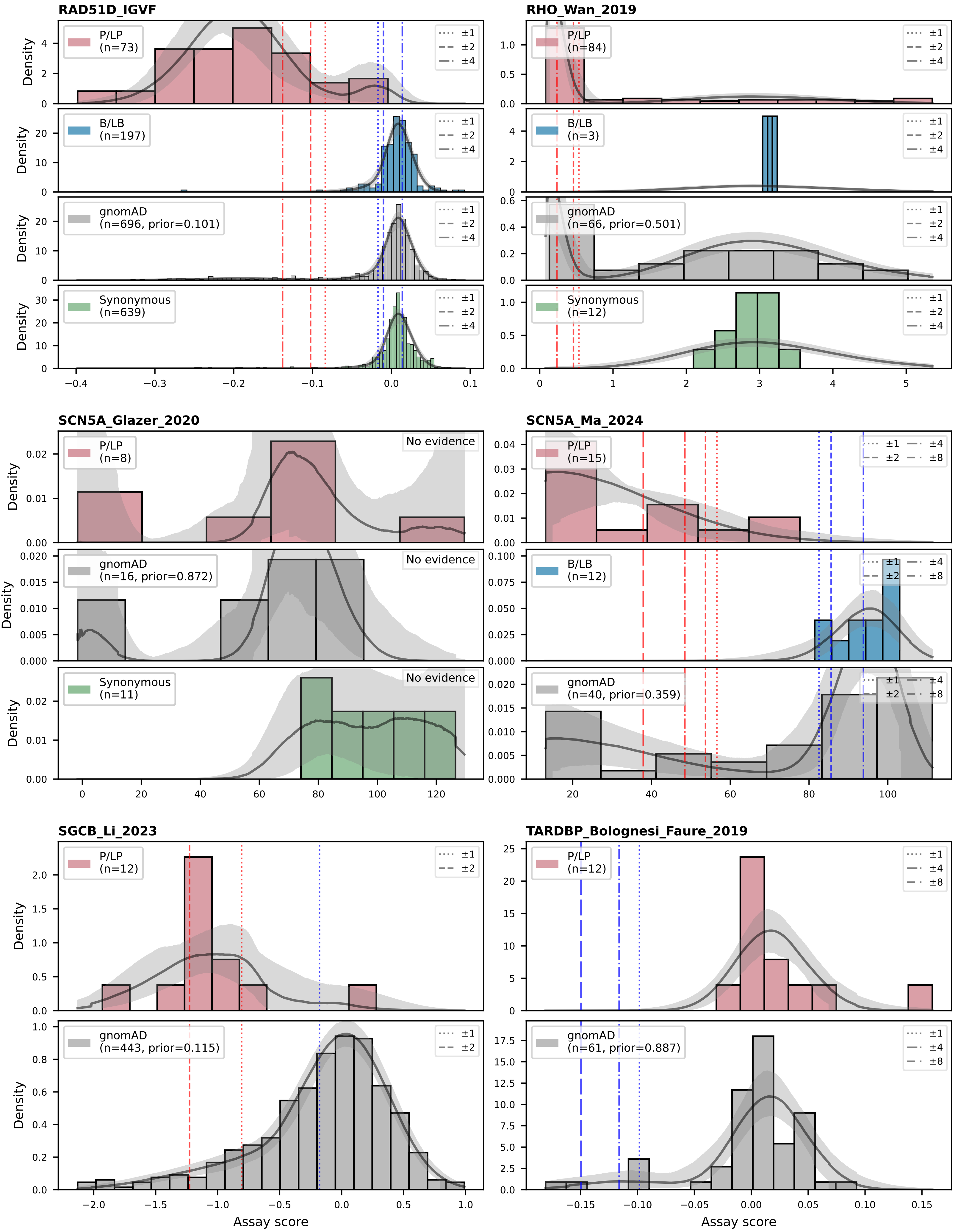

RAD51D_IGVF
P/LP (n=73)
B/LB (n=197)
gnomAD (n=696, prior=0.101)
Synonymous (n=639)
±1
±2
±4
Density
−0.4
−0.3
−0.2
−0.1
−0.0
0.1
RHO_Wan_2019
P/LP (n=84)
B/LB (n=3)
gnomAD (n=66, prior=0.501)
Synonymous (n=12)
0
1
2
3
4
5
SCN5A_Glazer_2020
P/LP (n=8)
gnomAD (n=16, prior=0.872)
Synonymous (n=11)
No evidence
0
20
40
60
80
100
120
SCN5A_Ma_2024
P/LP (n=15)
B/LB (n=12)
gnomAD (n=40, prior=0.359)
±8
SGCB_Li_2023
P/LP (n=12)
gnomAD (n=443, prior=0.115)
−2.0
−1.5
−1.0
−0.5
0.0
0.5
1.0
TARDBP_Bolognesi_Faure_2019
P/LP (n=12)
gnomAD (n=61, prior=0.887)
−0.15
−0.10
−0.05
0.00
0.05
0.10
0.15
Assay score

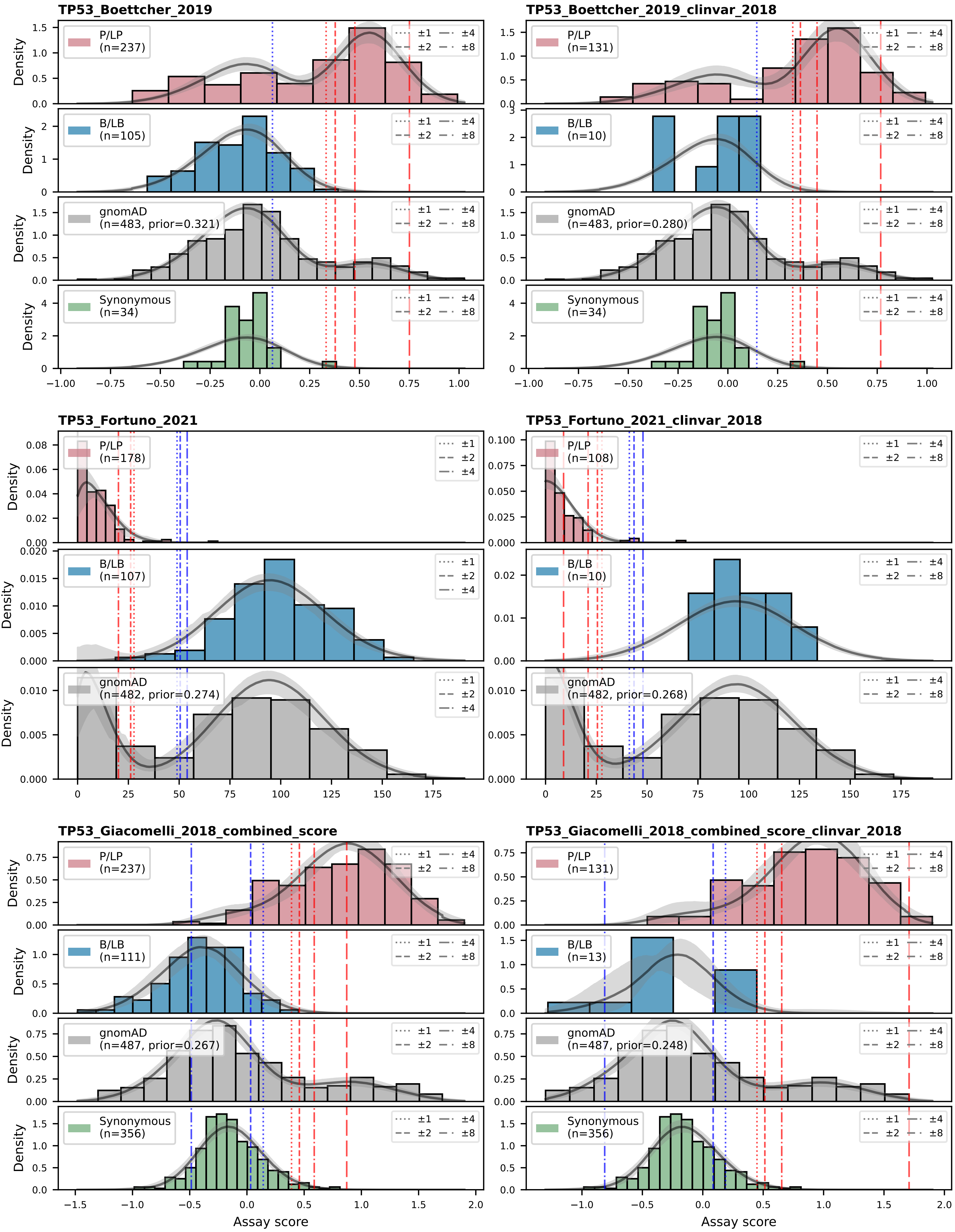

TP53_Boettcher_2019
TP53_Boettcher_2019_clinvar_2018
P/LP (n=237)
P/LP (n=131)
B/LB (n=105)
B/LB (n=10)
gnomAD (n=483, prior=0.321)
gnomAD (n=483, prior=0.280)
Synonymous (n=34)
±1 ±2 ±4 ±8
Density
TP53_Fortuno_2021
TP53_Fortuno_2021_clinvar_2018
P/LP (n=178)
P/LP (n=108)
B/LB (n=107)
B/LB (n=10)
gnomAD (n=482, prior=0.274)
gnomAD (n=482, prior=0.268)
TP53_Giacomelli_2018_combined_score
TP53_Giacomelli_2018_combined_score_clinvar_2018
P/LP (n=237)
P/LP (n=131)
B/LB (n=111)
B/LB (n=13)
gnomAD (n=487, prior=0.267)
gnomAD (n=487, prior=0.248)
Synonymous (n=356)
Assay score

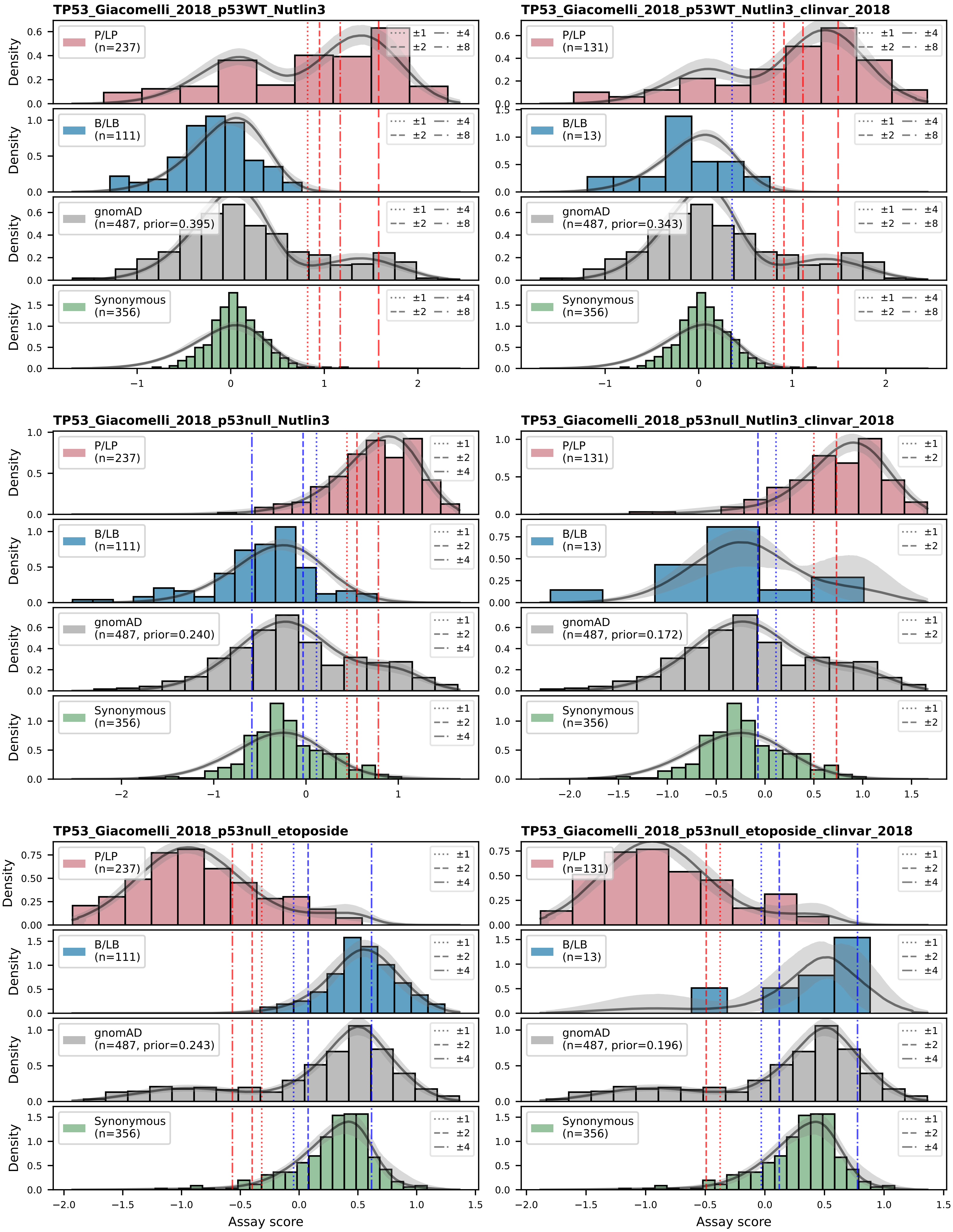
TP53_Giacomelli_2018_p53WT_Nutlin3
TP53_Giacomelli_2018_p53WT_Nutlin3_clinvar_2018
TP53_Giacomelli_2018_p53null_Nutlin3
TP53_Giacomelli_2018_p53null_Nutlin3_clinvar_2018
TP53_Giacomelli_2018_p53null_etoposide
TP53_Giacomelli_2018_p53null_etoposide_clinvar_2018
P/LP (n=237)
P/LP (n=131)
B/LB (n=111)
B/LB (n=13)
gnomAD (n=487, prior=0.395)
gnomAD (n=487, prior=0.343)
gnomAD (n=487, prior=0.240)
gnomAD (n=487, prior=0.172)
gnomAD (n=487, prior=0.243)
gnomAD (n=487, prior=0.196)
Synonymous (n=356)
±1 ±2 ±4 ±8
Density
Assay score

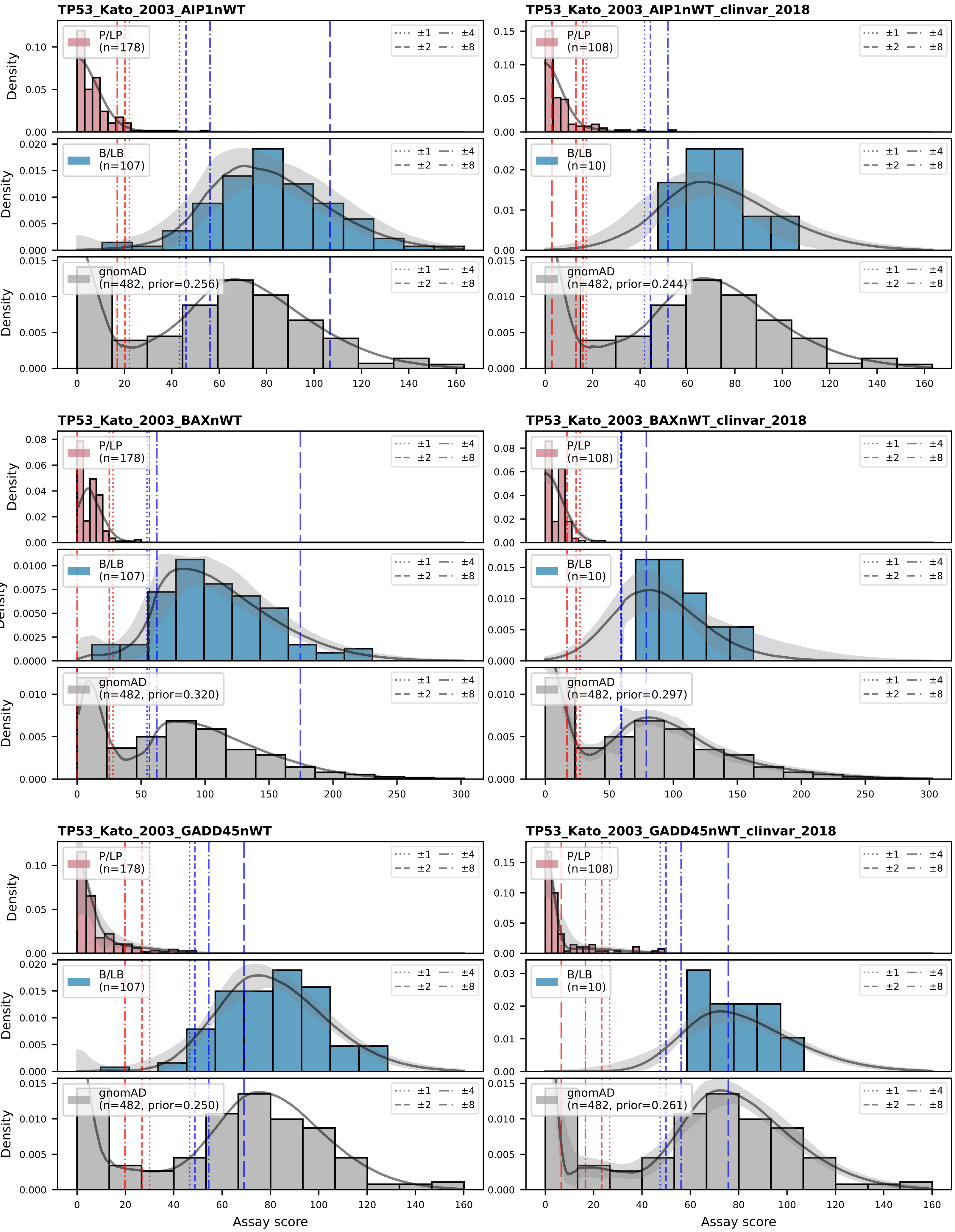
TP53_Kato_2003_AIP1nWT
TP53_Kato_2003_AIP1nWT_clinvar_2018
TP53_Kato_2003_BAXnWT
TP53_Kato_2003_BAXnWT_clinvar_2018
TP53_Kato_2003_GADD45nWT
TP53_Kato_2003_GADD45nWT_clinvar_2018
P/LP (n=178)
P/LP (n=108)
B/LB (n=107)
B/LB (n=10)
gnomAD (n=482, prior=0.256)
gnomAD (n=482, prior=0.244)
gnomAD (n=482, prior=0.320)
gnomAD (n=482, prior=0.297)
gnomAD (n=482, prior=0.250)
gnomAD (n=482, prior=0.261)
±1 ±2 ±4 ±8
Density
Assay score

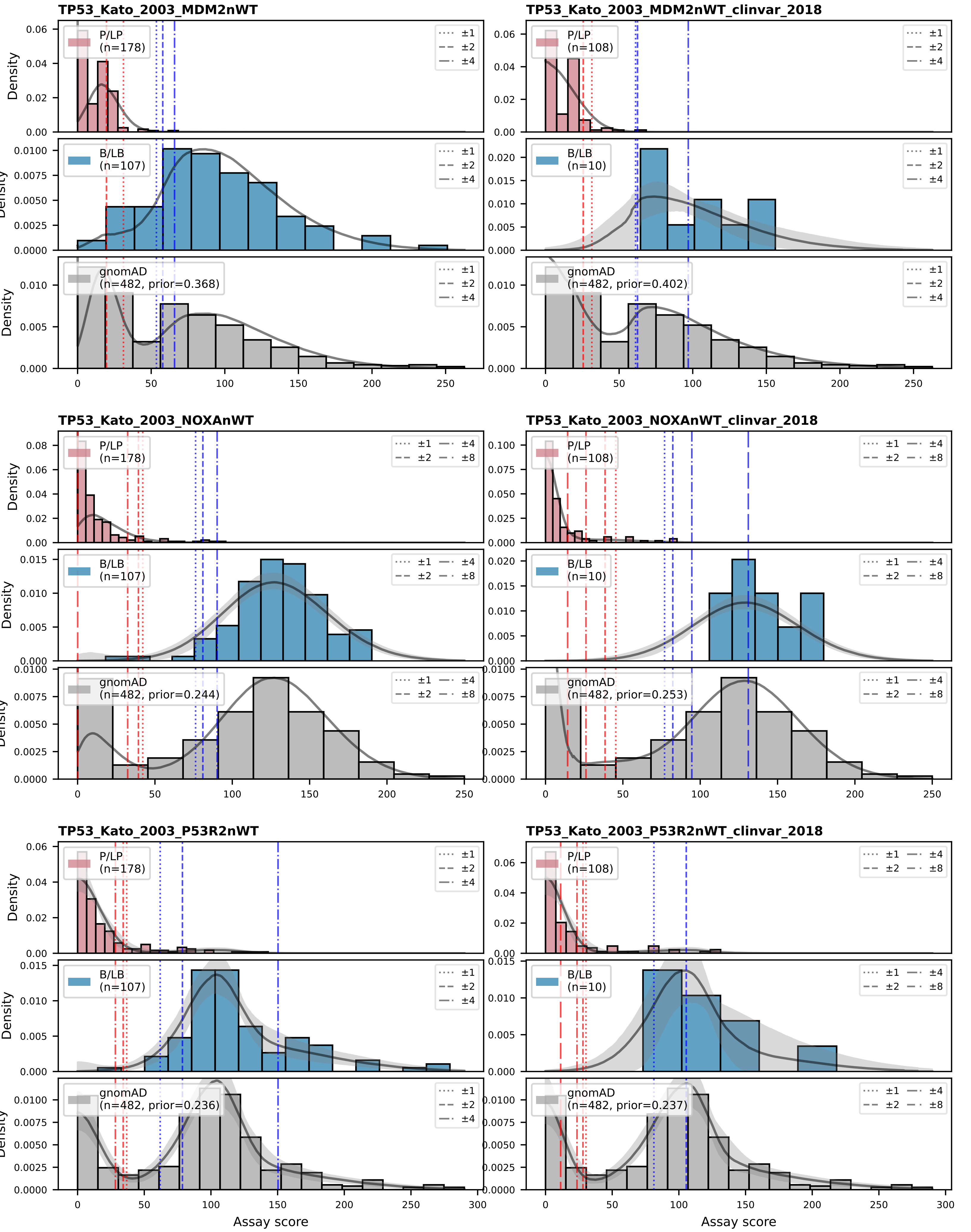
TP53_Kato_2003_MDM2nWT
TP53_Kato_2003_MDM2nWT_clinvar_2018
TP53_Kato_2003_NOXAnWT
TP53_Kato_2003_NOXAnWT_clinvar_2018
TP53_Kato_2003_P53R2nWT
TP53_Kato_2003_P53R2nWT_clinvar_2018
P/LP (n=178)
P/LP (n=108)
B/LB (n=107)
B/LB (n=10)
gnomAD (n=482, prior=0.368)
gnomAD (n=482, prior=0.402)
gnomAD (n=482, prior=0.244)
gnomAD (n=482, prior=0.253)
gnomAD (n=482, prior=0.236)
gnomAD (n=482, prior=0.237)
±1
±2
±4
±8
Density
Assay score

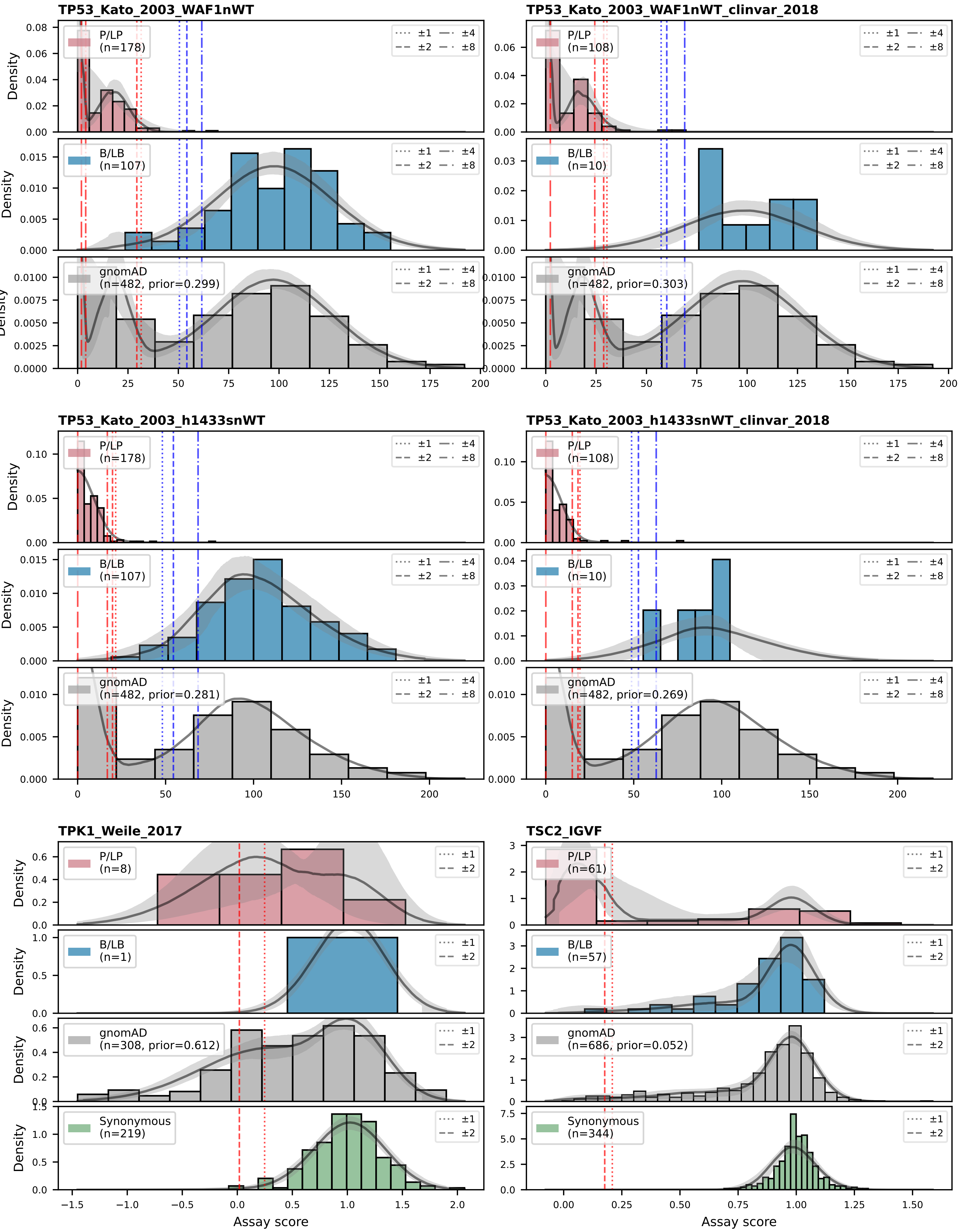
TP53_Kato_2003_WAF1nWT
TP53_Kato_2003_WAF1nWT_clinvar_2018
P/LP (n=178)
B/LB (n=107)
gnomAD (n=482, prior=0.299)
P/LP (n=108)
B/LB (n=10)
gnomAD (n=482, prior=0.303)
TP53_Kato_2003_h1433snWT
TP53_Kato_2003_h1433snWT_clinvar_2018
P/LP (n=178)
B/LB (n=107)
gnomAD (n=482, prior=0.281)
P/LP (n=108)
B/LB (n=10)
gnomAD (n=482, prior=0.269)
TPK1_Weile_2017
TSC2_IGVF
P/LP (n=8)
B/LB (n=1)
gnomAD (n=308, prior=0.612)
Synonymous (n=219)
P/LP (n=61)
B/LB (n=57)
gnomAD (n=686, prior=0.052)
Synonymous (n=344)
±1 ±2 ±4 ±8
Density
Assay score

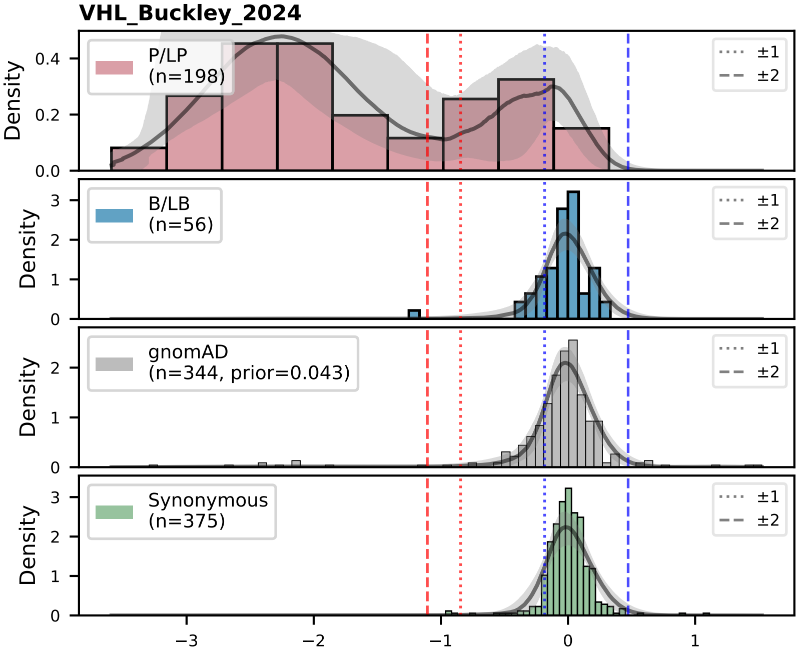
VHL_Buckley_2024
P/LP (n=198)
B/LB (n=56)
gnomAD (n=344, prior=0.043)
Synonymous (n=375)
±1
±2
Density

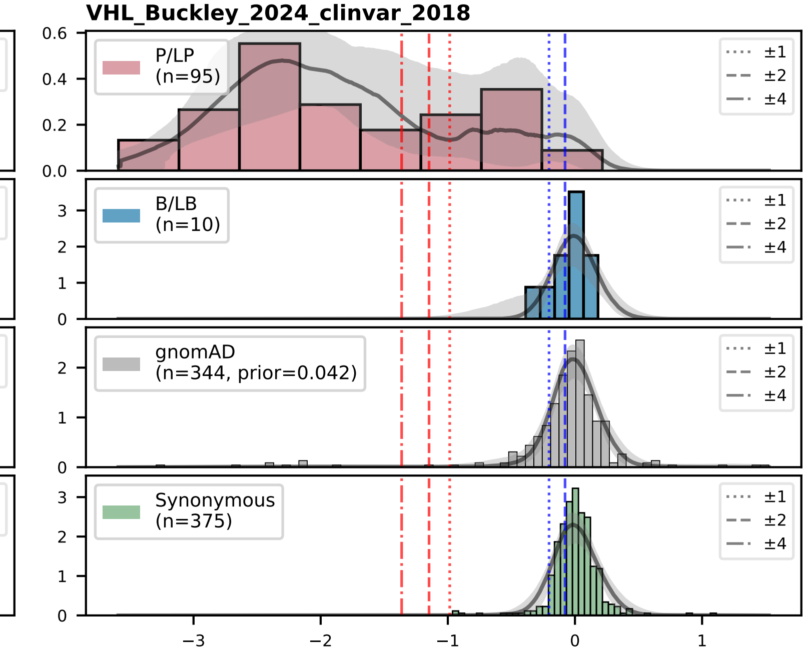
VHL_Buckley_2024_clinvar_2018
P/LP (n=95)
B/LB (n=10)
gnomAD (n=344, prior=0.042)
Synonymous (n=375)
±1
±2
±4

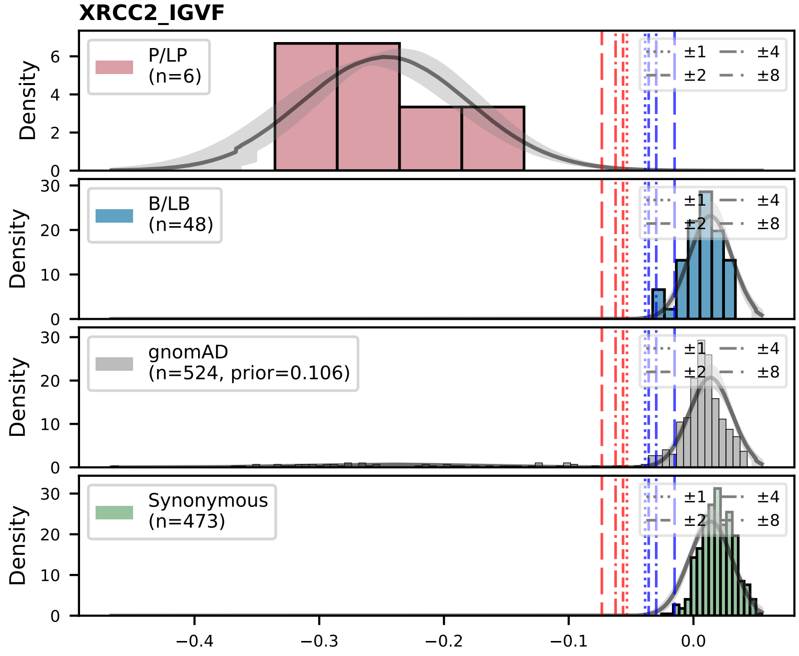
XRCC2_IGVF
P/LP (n=6)
B/LB (n=48)
gnomAD (n=524, prior=0.106)
Synonymous (n=473)
±1
±2
±4
±8
Density
